# RhoA GEF Mcf2lb regulates rosette integrity during collective cell migration

**DOI:** 10.1101/2023.04.19.537573

**Authors:** Hannah M. Olson, Amanda Maxfield, Nicholas L. Calistri, Laura M. Heiser, Alex V. Nechiporuk

## Abstract

During development, multicellular rosettes serve as important cellular intermediates in the formation of diverse organ systems. Multicellular rosettes are transient epithelial structures that are defined by the apical constriction of cells towards the rosette center. Due to the important role these structures play during development, understanding the molecular mechanisms by which rosettes are formed and maintained is of high interest. Utilizing the zebrafish posterior lateral line primordium (pLLP) as a model system, we identify the RhoA GEF Mcf2lb as a regulator of rosette integrity. The pLLP is a group of ∼150 cells that migrates along the zebrafish trunk and is organized into epithelial rosettes; these are deposited along the trunk and will differentiate into sensory organs called neuromasts (NMs). Using single-cell RNA sequencing and whole-mount in situ hybridization, we showed that *mcf2lb* is expressed in the pLLP during migration. Given the known role of RhoA in rosette formation, we asked whether Mcf2lb plays a role in regulating apical constriction of cells within rosettes. Live imaging and subsequent 3D analysis of *mcf2lb* mutant pLLP cells showed disrupted apical constriction and subsequent rosette organization. This in turn resulted in a unique posterior Lateral Line phenotype: an excess number of deposited NMs along the trunk of the zebrafish. Cell polarity markers ZO-1 and Par-3 were apically localized, indicating that pLLP cells are normally polarized. In contrast, signaling components that mediate apical constriction downstream of RhoA, Rock-2a and non-muscle Myosin II were diminished apically. Altogether our results suggest a model whereby Mcf2lb activates RhoA, which in turn activates downstream signaling machinery to induce and maintain apical constriction in cells incorporated into rosettes.

## INTRODUCTION

During development, cells undergo collective shape changes to accommodate organ morphogenesis. One prominent example of this behavior is in the formation of multicellular rosettes. Most multicellular rosettes are transient epithelial structures that contain five or more cells that interface at a central point, where apical membranes of these cells constrict. Multicellular rosettes are observed in many developmental contexts, including convergent extension during *Drosophila* embryogenesis, posterior Lateral Line (pLL) formation in zebrafish, vertebrate kidney tubule elongation, as well as numerous others (Blankenship et al., 2006; Gompel et al., 2001; Lienkamp et al., 2012).

Rosettes form through the process of apical constriction. Apical constriction occurs when the apical portion of an epithelial columnar cell narrows while the base of the cell remains at a constant width. This process is dependent on the contraction of the acto-myosin network (Nishimura and Takeichi, 2008; Ernst et al., 2012; Harding and Nechiporuk, 2012). In order for apical constriction to properly occur, cells first need to become polarized into apical and basal domains. The aPKC complex is a regulator of cell polarity and consists of aPKC, Par-6, and Par-3 (Chen and Macara, 2005). Proper activation and distribution of this complex ensures the apical localization of cell junction proteins including both adherens junction proteins (cadherin) and tight junction proteins (including ZO-1) (Niessen, 2007). Additionally, prior to constriction, the molecular players necessary for this process become apically localized, including F-actin fibers and non-muscle Myosin II. As rosettes are important intermediates during the formation of multiple organ systems, there has been a high interest in understanding molecular machinery that induces and maintains these structures.

To dissect this process, we use a well-established model system, the posterior lateral line primordium (pLLP). The pLLP is a group of ∼150 cells that is organized into polarized rosettes; each rosette gives rise to a sensory organ on the surface of the trunk called a neuromast (NM). NMs are part of the lateral line mechanosensory system that detects changes in water current and controls various swimming behaviors (Montgomery et al., 2000). During development, the pLLP forms just caudal to the otic vesicle. Between 22 and 48 hours post-fertilization (hpf), pLLP cells collectively migrate along the lateral aspect of the trunk (Metcalfe et al., 1985; Ghysen and Dambly-Chaudiere, 2004). Based on morphological and molecular differences, the migrating pLLP can be divided into two main regions, the leading region (leaders) and the trailing region (followers). Cells in the trailing region divide and differentiate during migration to form epithelial rosettes that are ultimately deposited during pLLP migration (Lecaudey et al., 2008; Nechiporuk and Raible, 2008). This results in the deposition of 5 to 6 NMs along the trunk. Cells in the leading region remain undifferentiated throughout most of migration and eventually form 2 to 3 NMs, called the terminal cluster at the distal end of the trunk (Aman and Piotrowski, 2008). The proximity of the pLLP to the skin makes it highly amenable to high-resolution live imaging. Together with the genetic tractability of zebrafish, this makes the pLLP an attractive model for studying mechanisms of rosette formation *in vivo*.

Two complimentary studies defined the molecular signaling pathways that regulate rosette formation in both the leading and trailing cells of the pLLP (Ernst et al., 2012; Harding and Nechiporuk, 2012). Harding and Nechiporuk (2012) identified the signaling pathway in the leading portion of the pLLP that dictates apical constriction and rosette formation (Harding and Nechiporuk, 2012). In this study, researchers found that Fgf ligands signal through the Fgf receptor to activate Ras-MAPK signaling to induce apical localization of the Rho kinase Rock-2a (Harding and Nechiporuk, 2012). RhoA activation of Rock-2a induces phosphorylation of non-muscle Myosin II, initiating the constriction of apically localized actin fibers (Harding and Nechiporuk, 2012). In a parallel study, Ernst et al. (2012) identified the scaffolding protein Schroom3 as a transcriptional target of Fgf signaling within the trailing region of the pLLP (Ernst et al., 2012). The authors showed that knockdown of Schroom3 results in impaired apical constriction and rosette formation, a phenotype similarly observed with Fgf inhibition (Ernst et al., 2012). Similar to the leading cells, activation of Rock-2a and subsequent phosphorylation of non-muscle Myosin II in the trailing rosettes are necessary for apical constriction and rosette formation. These two studies show that the signaling mechanisms to induce apical constriction both in the leading and the trailing region are similar and involve activation of RhoA and its downstream effectors. However, how RhoA activity is regulated and maintained in this context is not known.

In this study, we identified Mcf2lb as a regulator of rosette integrity during pLLP migration. We showed that loss of *mcf2lb* results in an excess number of deposited NMs along the trunk of the embryo. Using live imaging, we showed that this phenotype results from deposition of rosette clusters in the mutant, instead of evenly spaced single rosettes in WT. We further demonstrated that this behavior results from abnormal rosette organization in the *mcf2lb* mutant pLLP. 3D analysis of individual mutant cells revealed impaired apical constriction. Notably, the tight junction marker ZO-1 and polarity marker Par-3 remained properly polarized, indicating that rosette cells are properly polarized. In contrast, immunostaining of downstream RhoA signaling components, Rock-2a and the non-muscle Myosin II component Myosin Regulatory Light Chain (MRLC), showed diminished signal at the rosette centers in *mcf2lb* mutants. These results suggest a model by which Mcf2lb acts as a GEF to activate RhoA, which then can induce RhoA signaling components to initiate apical constriction. In this study, we identified a novel regulator of apical constriction and rosette integrity and demonstrated that maintenance of proper rosette integrity is necessary in the formation of the sensory system, the pLL.

## MATERIALS AND METHODS

### Zebrafish husbandry and strains

Adult zebrafish were maintained under standard conditions (Westerfield, 2000). Membranes of the cells in the pLLP were visualized using Tg(*-8.0claudinB:lynGFP*)*^zf106^* and Tg(*prim:lyn2-mCherry*) (Haas and Gilmour, 2006). Tg(*prim:lyn2-mCherry*) is a fortuitous integration of the memRFP, driven by a 3 kb *sox10* promoter that in addition to the neural crest also labels the pLLP (Wang et al., 2018). TgBAC(*cxcr4b:F-tractin-mCherry*) was used to label the pLLP in red fluorescence for single-cell RNA sequencing (Yamaguchi et al., 2022). TgBAC(*cxcr4b; LexPr,cryaa:GFP)* was used in combination with the plasmid pDest-Cg2-LexOP-secGFP to mosaically induce expression of sec-GFP in the pLLP (Durdu et al, 2014).

### Embryo dissociation and FACS

30 hpf Tg(*-8.0claudinB:lynGFP*)*^zf106^*/ TgBAC(*cxcr4b:F-tractin-mCherry*) zebrafish embryos were collected and euthanized in 1.7 ml microcentrifuge tubes. Embryos were deyolked using a calcium-free Ringer’s solution (116 mM NaCl, 2.6 mM KCl, 5 mM HEPES pH 7.0), by gently pipetting up and down with a P200 pipet. Embryos were incubated for 5 minutes in Ringer’s solution. Embryos were transferred to pre-warmed protease solutions (0.25% trypsin, 1 mM EDTA, pH 8.0, PBS) and collagenase P/HBSS (100 mg/mL) was added. Embryos were incubated at 28° C for 15 minutes and were homogenized every 5 minutes using a P1000 pipet. The Stop solution (6X, 30% calf serum, 6 mM CaCl_2_, PBS) was added and samples were centrifuged (350xg, 4° C for 5 minutes). Supernatant was removed and 1 mL of chilled suspension solution was added (1% FBS, 0.8 mM CaCl_2_, 50 U/mL penicillin, 0.05 mg/mL streptomycin, DMEM). Samples were centrifuged again (350g, 4° C for 5 minutes) and supernatant was removed. 700 μl of chilled suspension solution was added and cells were resuspended by pipetting. Cells were passed through a 40 μm cell strainer into a FACs tube and kept on ice. GFP and RFP+ cells were FAC sorted on a BD Symphony cell sorter into sorting buffer (50 μl PBS/ 2% BSA) in a siliconized 1.5mL tube.

### 10X chromium scRNA-seq library construction

FAC-sorted live cells were used for scRNA-seq. Approximately 15,000 cells were loaded into the Chromium Single Cell Controller to generate barcoded RT products (Chemistry Ver 3.0; 10X Genomics, Pleasanton, CA. USA). The library was sequenced using Illumina NovaSeq 6000 to a depth of at least 50,000 reads per cell.

### Quality Control and Unsupervised clustering

Single cell reads were aligned to Ensembl version GRCz11 of the zebrafish genome by the Integrated Genomics Laboratory (Oregon Health & Science University) using Cell Ranger (version 3.1.0; 10X Genomics, Pleasanton, CA. USA). UMI count matrix was analyzed using Seurat (version 4.0.1) (Butler et al., 2018). Quality control filtered out genes expressed in less than three cells, cells with less than 1,900 unique genes, and cells that expressed greater than 5% of mitochondrial transcripts. The remaining 3,851 cells subjected to further analysis. Linear Dimensionality reduction, clustering and UMAP visualization were performed with Seurat (Butler et al., 2018). Briefly, Principal Component analysis was performed to project the 2,000 most variable genes into 20 principal components. Twenty-two clusters were identified with the Seurat implementation of the Louvain Algorithm using a resolution of 0.8, and visualized with 2 UMAP dimensions. Clusters were manually annotated using a whole zebrafish single-cell transcriptome atlas (Farnsworth et al., 2020) as well zebrafish database of gene expression (ZFIN expression atlas: Thisse et al., 2001). Notably, a cluster optimization algorithm (Lun, 2022) identified resolution 0.7 as optimal. However in that case, *msx1b*+ fin epidermal cells co-clustered with pLLP cells; these populations separated into distinct clusters at resolution 0.8, which we used for further analysis. A subset of pLLP clusters were identified using literature derived markers, and then subclustered to identify unique pLLP transcriptomic states (Farnsworth et al., 2020). pLLP cells subclustered into 3 groups using resolution 0.2. Although, cluster optimization identified resolution 0.3 as optimal, leading to four clusters. This was a result of the follower cells separating into two, very similar populations (Supplemental Fig. 6). As such, we used resolution 0.2 for further analysis.

### Gene Ontology analysis

Gene Ontology (GO) analysis was performed using the following R packages GO.db (Carlson, 2019), biomaRt (Durnick et al., 2005), clusterProfiler (Yu et al., 2012), and org.Dr.eg.db (Carlson, 2019). KEGG pathway analysis was performed using the DAVID online platform (Dennis et al., 2003; Hosack et al., 2003).

### In situ hybridization and whole mount immunostaining

RNA in situ hybridization was performed as described previously (Andermann et al., 2002). Digoxygenin-labeled antisense RNA probes were generated for the following genes: *mcf2lb*, *twf2b*, *arhgef4*, *twf2b*, *atoh1a* (Itoh and Chitnis 2001), and *deltaA* (Itoh and Chitnis 2001).

Forty-five hpf embryos were collected and fixed in BT fixative (anti-ZO-1), glyofixx (Thermo Scientific, anti-Rock2a), or Bouin’s fixative (Polysciences; anti-pMRLC) (Westerfield, 2000b) overnight at 4° C. After removing the fixative, embryos were washed with PBS/0.1% Triton washes, embryos were blocked with PBTx/5% goat serum/1% bovin serum albumin, 1% DMSO before incubating in primary antibody at 4° C overnight. The embryos were washed in PBS/0.1% Triton and incubated in secondary antibody at 4°C overnight and then washed with PBS/0.1% Triton. After the final wash, embryos were mounted in 50% PBS/50% glycerol for imaging. Primary antibodies were used at the following dilutions: mouse anti-ZO-1 (1:500), rabbit anti-Rock2a (1:50), and rabbit anti-pMRLC (1:20). Secondary antibodies (used at 1:750 concentration) were goat anti-rabbit Alexa Fluor 568, goat anti-mouse Alexa Fluor 568 and goat anti-chicken Alexa Fluor 488 (Thermo Fisher Scientific). Nuclei were visualized with DAPI.

### CRISPR-Cas9-mediated knockout

Three single guide RNAs targeting exon 6, 10, and 21 for CRISPR-Cas9 targeting of *mcf2lb* were designed and injected as previously described (Shah et al., 2015) (Key resources table 1). All 3 guides were injected together into Tg(*8.0claudinB:lynGFP*)*^zf106^* – positive embryos. At 3 dpf, embryos were screened for a pLL formation phenotype and genotyped to assess CRISPR efficiency. F0 - injected embryos were raised to adulthood, were in-crossed, and F1 embryos were screened at 3 dpf for a pLL formation phenotype. Positive F0 adults were out-crossed to a WT background, and progeny were raised to adulthood. A stable line was identified when F1 adults were in-crossed and their subsequent progeny were screened at 3 dpf for the pLL phenotype, which was observed with Mendelian inheritance.

**Table 1:**
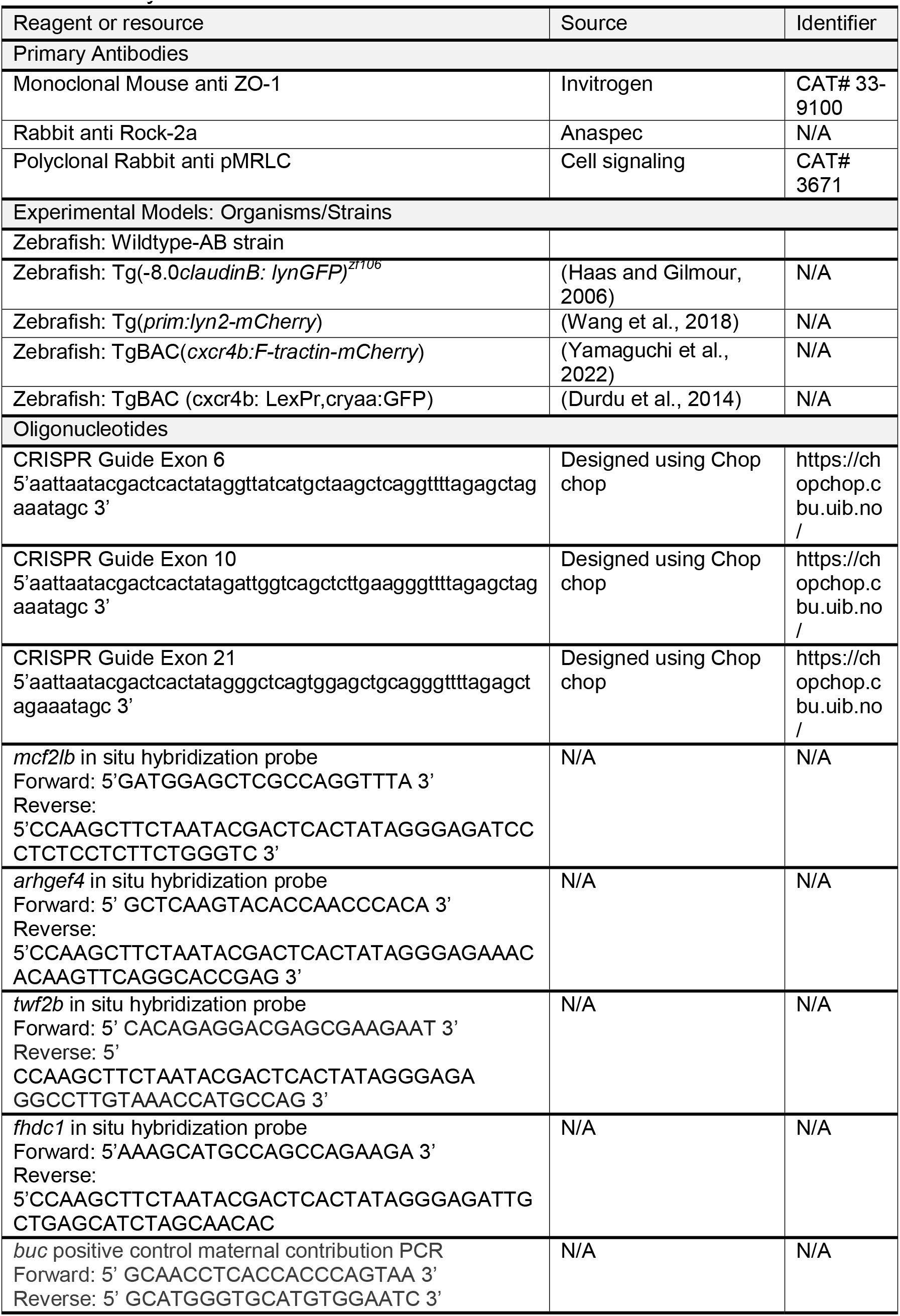
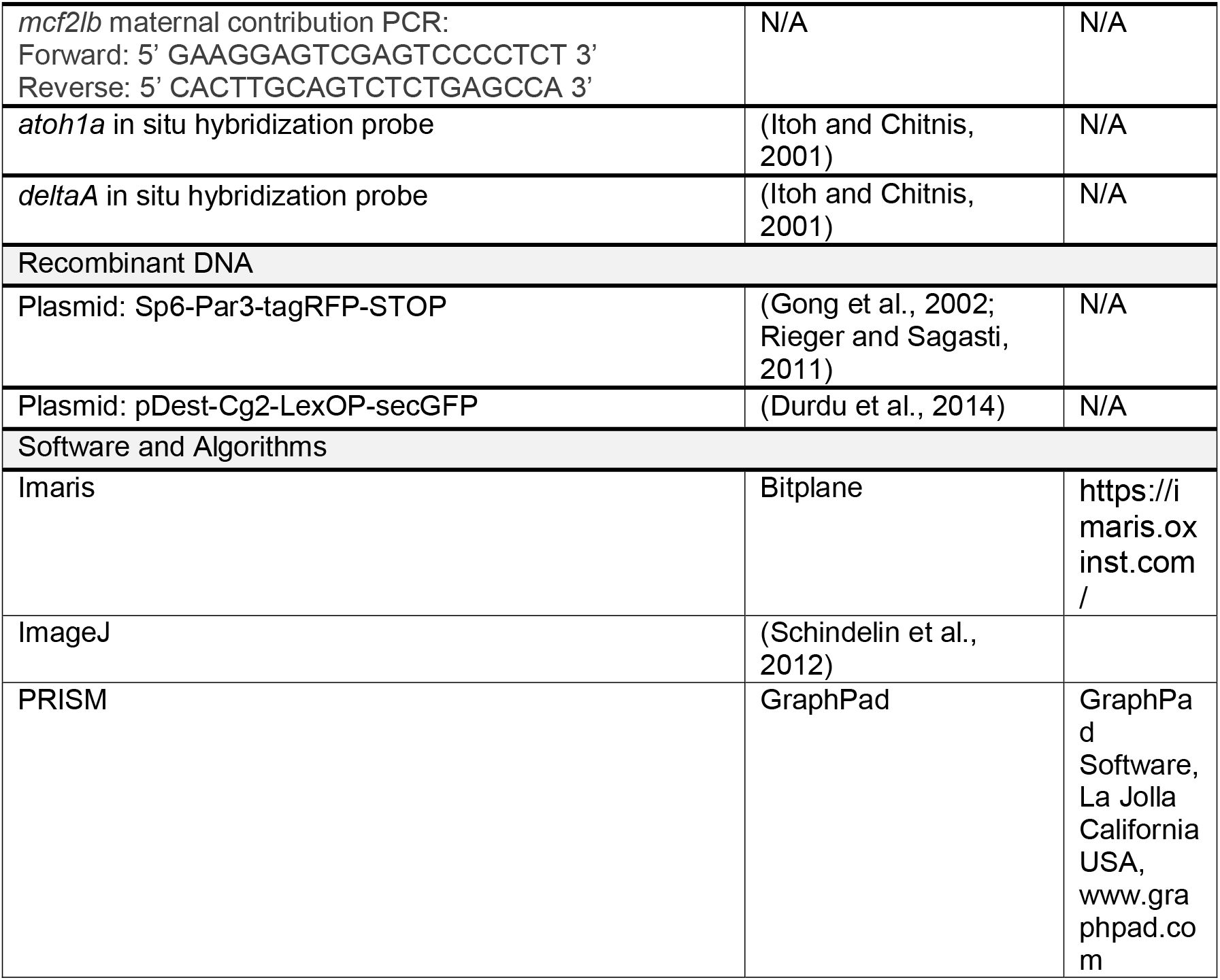
Key Resources Table.

All three guides were efficient in their cutting and produced indels. We generated two mutant alleles that contained mutations in three distinct exons (Supplemental Fig. 3A). The first allele, nl25 – allele 1, contained an 8 bp insertion in exon 6 which produces an early STOP within the insertion; a 6 bp deletion and an 18 bp insertion in exon 10 (a net 12 bp insertion) that also ultimately leads to an early STOP. The second allele, nl26 – allele 2, contained a 7 bp insertion in exon 6 and results in an early STOP shortly after the insertion; a 442 bp deletion and 18 bp insertion that produces a net 424 bp deletion. In addition, both alleles contained a 7 bp deletion in exon 21 (early STOP shortly after the deletion) (Supplemental Fig. 3A). The *mcf2lb* mutant population used in this study contains a mixed population of both nl25 – allele 1 and nl26 – allele 2. We observed no phenotypic differences between the two alleles (Supplemental Fig. 3B).

To distinguish between nl25 and nl26, regions including exon 6, exon 10, and exon 21 were PCR amplified from adult genomic DNA and digested with restriction enzymes. Phusion High-Fidelity DNA Polymerase (ThermoFisher) was used for exon 6, and Taq DNA Polymerase (NEB) was used for exon 10 and 21. Standard PCR conditions according to the manufacturers’ instructions were used. Annealing temperatures of 63°C, 62°C, and 55°C were used for exon 6, exon 10, and exon 21 PCR amplification, respectively. PCR-amplified DNA was digested with MwoI and HpyCH4III in separate reactions for exon 6, DrdI for exon 10, and Cac8I for exon 21. Amplification of the region of DNA containing exon 10 in nl26 proved to be challenging.

### Plasmids and injections and LexPr/LexOP induction

Sp6-Par3-tagRFP plasmid was used from (Harding and Nechiporuk, 2012). *par3-tagRFP* mRNA was synthesized using the mMessage mMachine Kit (Life Technologies) and microinjected at 250 pg/embryo. pDEST-Cg2-LexOP-secGFP (Wang et al., 2018) was microinjected into *TgBAC(cxcr4b:LexPr,cryaaGFP)* transgenic embryos (Durdu et al., 2014) at 5 pg/embryo. Expression of LexOP-secGFP was induced by treating 29 hpf TgBAC(*cxcr4b:LexPr; cryaa:GFP*) transgenic embryos with 20 μM mifepristone (RU486, Sigma) for 6 hours at 31°C.

### Transplantation experiments and time-lapse live imaging

Transplantation experiments were carried out as previously described (Nechiporuk and Raible, 2008). All host embryos expressed the Tg(*-8.0claudinB:lynGFP*)*^zf106^*transgene and were either in a WT or *mcf2lb* mutant background. Donor cells were derived from WT or *mcf2lb* mutant Tg(*prim:lyn2-mCherry*) transgenic embryos. Embryos were screened at ∼28 hpf and then mounted at ∼30 hpf for live imaging.

For time-lapse imaging, embryos were anesthetized in 0.02% tricaine (MS-222; Sigma) embedded in 1.2% low-melting point agarose and imaged either using a 20X/NA = 0.95 water dipping lens or a 40X/ NA = 1.25 silicone lens on an upright Fluoview 3000 confocal microscope (Olympus). For overnight time-lapse imaging, pLLPs were imaged between 30-44 hpf. For high-resolution pLLP imaging and imaging of Par-3-tagRFP in the migrating pLLP, pLLPs were imaged ∼30 hpf.

### Foci vs. points distinction

Gatherings of membranes were depicted as either foci or points. To determine what qualifies as a focus vs. a point, the width and height of each gathering of membrane in WT pLLP was measured. These values were averaged and a gathering of membrane was determined as a focus if its value landed between one standard deviation above or below the average of the width <u>and</u> the height of all WT foci. If a gathering of membrane did not meet these criteria, it was determined to be a point.

### Apical constriction quantification

Using the Imaris Cells function, all cells in the pLLP were 3D reconstructed. In the trailing 70% of the pLLP, all the Apical Constriction Index’s (ACI) of all cells were measured. ACIs are the ratio between the apical width of the cell, 1 μm below the apical top of the cell, and the basal width, 1 μm above the basal bottom of the cell. Cells were categorized as cells incorporated into rosettes if they touched or reached towards a focus or point. Cells were categorized as touching multiple points if cells contacted multiple foci or points. Cells were not included in analysis if they were dividing, did not span the entire length of the pLLP, or were considered sheath cells, those cells that lie entirely flat along the top of the pLLP, closest to the skin. All other cells were categorized as not being incorporated into the rosettes.

### Membrane variability and secGFP quantification

The apical width 1 μm below the top of the transplanted cell was measured throughout a 1-hour imaging period. Membrane variability was defined as the standard deviation of the apical membrane width of a cell over the course of this imaging period. Normalized fluorescence intensity of secGFP was measured by dividing the total fluorescence intensity of GFP in the microlumen by the total fluorescence intensity of cells expressing secGFP that contribute to the rosette centers of trailing protoNMs of pLLPs and deposited NM2. The same circular ROI was used to measure the fluorescence intensity in the presumptive microlumen across all samples.

### Polarity marker and RhoA downstream signaling component quantification

Fluorescence intensity values for ZO-1 and Par3 were obtained by measuring the fluorescence intensity throughout the entire pLLP. For ZO-1, the fluorescence intensity of an ROI for the top half of the pLLP was measured. For Par-3 the fluorescence intensity of an ROI that was the entire length of the pLLP, half of its width and half of its height was measured. These ROI values were then divided by the sum value in the pLLP to achieve the percentage of fluorescence intensity that was apically localized or the percentage of fluorescence intensity that was midline and apically localized (respectively).

For ZO-1, NMs values were obtained by measuring the fluorescence intensity throughout the whole NM and measuring the fluorescence intensity in an ROI that was the top half of the NM. The ROI value was then divided by the sum value in the pLLP to achieve the percentage of apically localized signal.

For Rock-2a and pMRLC quantification, fluorescent intensity values were obtained by measuring fluorescence intensity throughout the entire pLLP and measuring the fluorescence intensity in an ROI around the rosette centers. ROI was a consistent size across Rock-2a analyzed pLLPs and pMRLC analyzed pLLPs. ROI was determined by using the average width and length of a focus and one-third of the average depth of the pLLP.

### Image Processing

Images were processed using ImageJ (Abramoff et al., 2004; Schindelin et al., 2012) or Imaris (Bitplane) software.

### Statistics

Data were analyzed in PRISM and R. Data were suitable for parametric analysis unless otherwise noted. A Mann-Whitney U assuming equal variances was used to compare ACIs of cells incorporated into rosettes in WT and *mcf2lb* mutant pLLPs, to compare apical width in WT and *mcf2lb* mutant pLLPs, to compare basal width in WT and *mcf2lb* mutant pLLPs, and to compare membrane variability in WT and *mcf2lb* mutant transplanted cells.

### Data Access

scRNA-seq data is publicly available through Gene Expression Omnibus (GEO) database, accession# GSE229567. The code used for scRNA-seq data analyses as well as to generate panel (D) in Fig. 6 is available on Github: https://github.com/anechipor/nechiporuk-lab-Olson_et_al_2023.

## RESULTS

### Identification of factors that regulate actin dynamics in the pLLP

Due to the important role of the actin cytoskeleton in both pLLP protrusive behavior as well as in the morphological changes that occur during rosette formation within the pLLP, we set out to identify potential genetic regulators of actin dynamics (Ernst et al., 2012; Harding and Nechiporuk, 2012; Dalle Nogare et al., 2020; Olson and Nechiporuk, 2021; Yamaguchi et al., 2022). To achieve this, we first identified the transcriptional profile of the pLLP by performing single-cell RNA sequencing (scRNA-seq). We used fluorescence-activated cell sorting (FACs) to isolate cells from 30 hpf zebrafish embryos that carried two transgenes marking the pLLP, Tg(-8.0*claudinB: lynGFP)^zf106^* (Haas and Gilmour, 2006) and TgBAC(*cxcr4b:F-tractin-mCherry)^p3^*(Yamaguchi et al., 2022). GFP+ and mCherry+ cells were processed using the 10X Chromium platform, and libraries were sequenced at approximately 50,000 reads per cell. scRNA-seq data were subjected to a quality control and unsupervised clustering using the Seurat pipeline (Butler et al., 2018). Unsupervised clustering and UMAP reduction resulted in 22 individual clusters that are annotated in Fig. 1A. pLLP cells were identified as clusters expressing known lateral line markers, including *hmx2* and *hmx3a* (Fig. 1A-C). Reclustering of the subset of pLLP cells did not reveal any additional transcriptional heterogeneity with three pLLP clusters, although distribution of some cells in clusters 0 and 1 changed (Fig. 1D).

**Figure 1:**
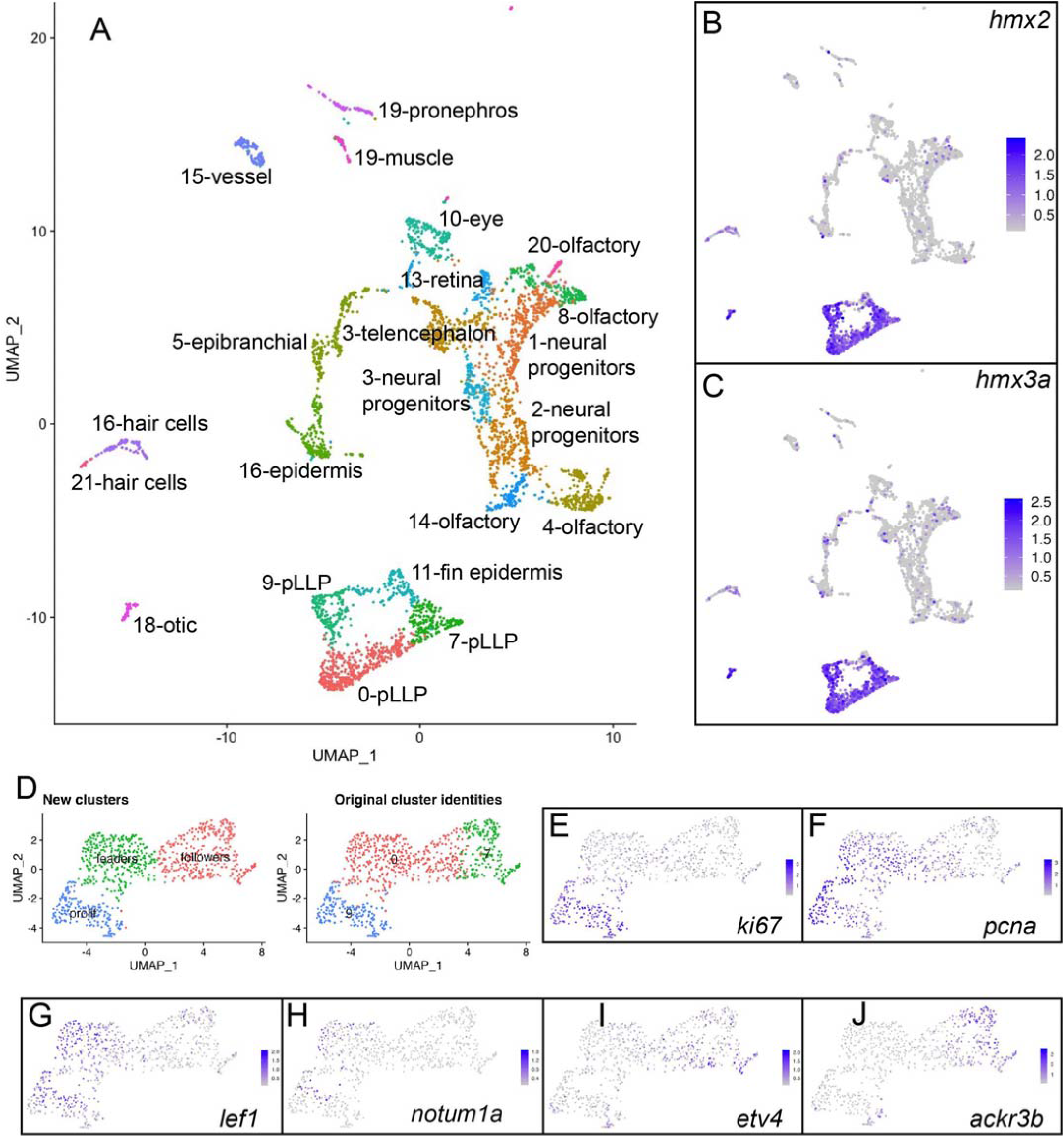
Identification of the pLLP transcriptional profile by scRNA-seq. (A) Unsupervised clustering and UMAP reduction diagram of cells derived from Tg(-8.0*claudinB: lynGFP)^zf106^*; TgBAC(*cxcr4b:F-tractin-mCherry)^p3^*transgenic embryos. (B, C) Feature plots of *hmx2* and *hmx3a* identify clusters 0, 7, and 9 as pLLP. (D) Unsupervised reclustering of clusters 0, 7, and 9. (E, F) Feature plots of *ki67* and *pcna* identify cluster 2 as proliferating cells. (G, H) Feature plots of *lef1* and *notum1a* identify cluster 1 as leader cells. (I, J) Feature plots of *etv4* and *ackr3b* (also known as *cxcr7b*) identify cluster 0 as follower cells.

Proliferation markers *ki67* and *pcna* identified cluster 2 as proliferating pLLP cells (Fig. 1E, F) (Gerdes et al., 1984; Celis and Celis, 1985). Previous studies showed that the Wnt signaling pathway is active in leading cells (Aman and Piotrowski, 2008). In contrast, downstream Fgf signaling components and the chemokine receptor *ackr3b* (also known as *cxcr7b*) are expressed in trailing pLLP cells (Raible and Brand, 2001; Dambly-Chaudiere et al., 2007). *lef1* and *notum1a*, two Wnt pathway signaling components (Giraldez et al., 2002; Clevers, 2006), were upregulated in cluster 1, whereas *etv4* and *ackr3b* were upregulated in cluster 0 (Fig 1G-J). Thus, our three subclusters mark transcriptionally distinct leader, follower, and proliferating pLLP cells.

We next investigated expression of genes that regulate actin dynamics in pLLP cells. To achieve this, we used Seurat function AddModuleScore to create a gene signature for GO (Gene Ontology) terms associated with actin regulation and actin dynamics (Ashburner et al., 2000). We then applied clusterProfiler package to analyze these gene signatures in pLLP cells (Gene Ontology Consortium). This analysis showed that most of the GO term signatures were enriched in pLLP cells. (Supplemental Fig. 1A). Enrichment of components that regulate actin dynamics is also illustrated on the Regulation of Actin Cytoskeleton KEGG (Kyoto Encyclopedia of Genes and Genomes) pathway map (Supplemental Fig. 1B) (Kanehisa et at., 2000). To visualize expression of individual genes, we grouped GO terms into: 1) actin binding; 2) Rho-GTPases, GAPs, GEFs; and 3) actin polymerization categories (Supplemental Fig. 2). These analyses demonstrated that a vast majority of actin regulatory genes have higher levels of expression in the follower cell cluster in comparison to either the leader or the proliferating cell cluster (Supplemental Fig. 1A and 2).

**Figure 2:**
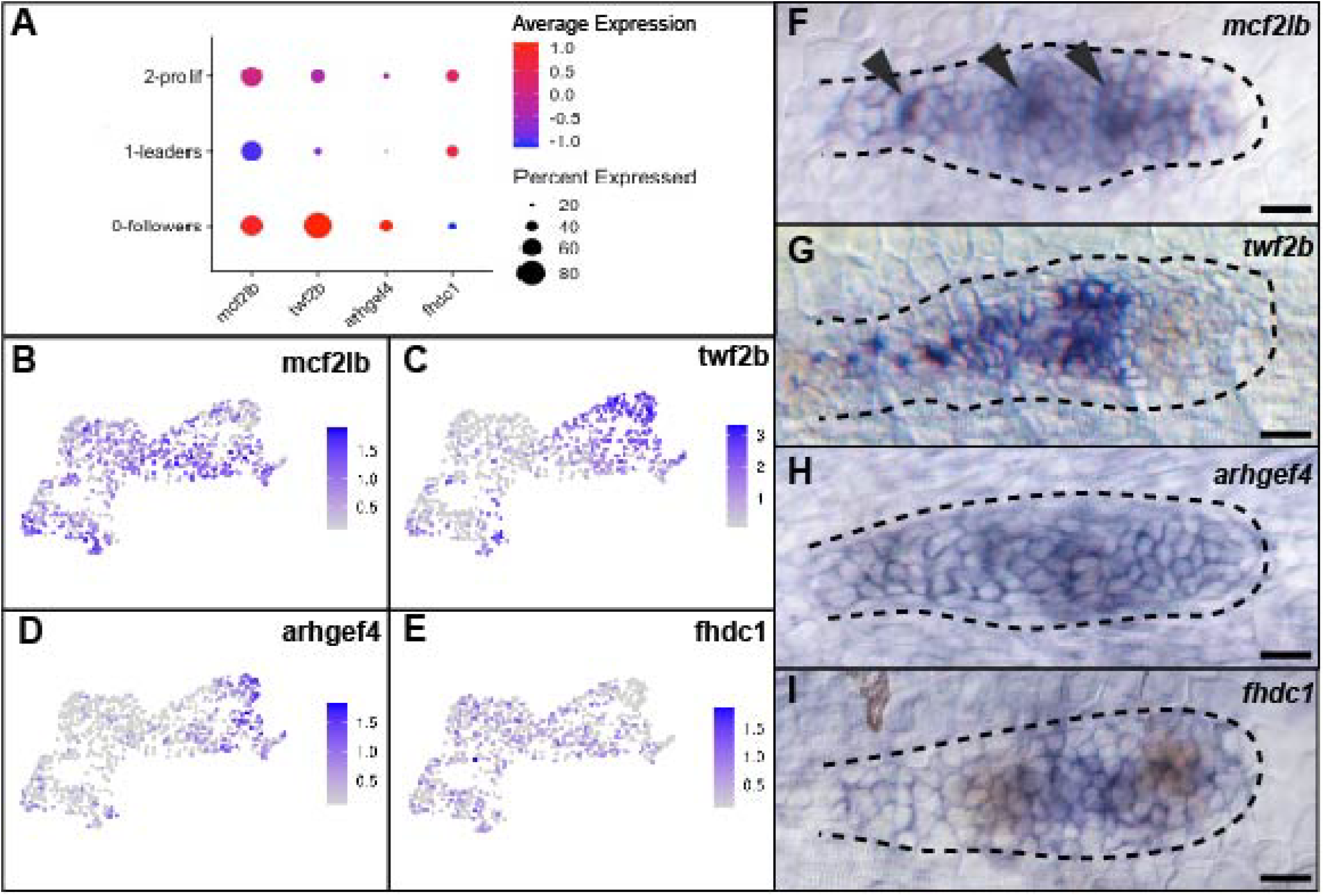
Expression profile of genes that regulate actin dynamics. (A) Dot plot expression profile of *mcf2lb*, *twf2b*, *arhgef4*, and *fhdc1*. (B – E) Feature plots showing expression profile of the above four genes in pLLP clusters. (F – I) In situ hybridization of the four genes. Note that expression profiles via in situ hybridization largely match those observed by scRNA-seq. Arrowheads indicate rosette centers. Dotted lines indicate the outline of the pLLP. Scale bars for panels F – I = 10 μm.

### scRNA-seq validation by in situ hybridization

To validate our scRNA-seq dataset, we used in situ hybridization to visualize expression of four genes, *mcf2lb*, *twf2b*, *arghef4*, and *fhdc1* in the pLLP. We chose these genes as they regulate different aspects of actin dynamics and show region-specific expression (Fig. 2). *mcf2lb* encodes a known GEF for RhoA, Rac1, and Cdc42 in different contexts with the majority of the evidence supporting its role as a GEF for RhoA (Horii et al., 1994; Whitehead et al., 1999; Cheng et al., 2004). In situ hybridization for *mcf2lb* was consistent with the scRNA-seq: it is expressed throughout the pLLP, with lower levels in the leading region (Fig. 2 A, B, F). Interestingly, the *mcf2lb* transcript appears to be enriched in the cells that form pLLP rosettes (Fig. 2F, arrowheads). *twf2b* encodes an f-actin capping protein that sequesters g-actin, thus inhibiting actin polymerization; it is expressed almost exclusively in the follower cells by both scRNA-seq and in situ (Fig. 2 A, C, G) (Vartiainen et al., 2003; Nevalainen et al., 2009). *arghef4* encodes a GEF for RhoA, Rac1, and Cdc42 and is upregulated in the followers of the pLLP via scRNA-seq. In situ hybridization revealed that *arghef4* is mostly expressed throughout the pLLP with some higher levels in the follower cells (Fig. 2 A, D, H) (Kawasaki et al., 2000; Gotthardt and Ahmadian, 2007). Finally, *fhdc1* encodes an actin-binding protein involved in stress fiber formation; it is expressed higher in the leading cells by in situ, which is consistent with the scRNA-seq (Fig. 2 A, E, I) (Young et al., 2008). In summary, our *in vivo* expression patterns by in situ largely confirm those observed by scRNA-seq.

### Loss of *mcf2lb* results in the supernumerary deposition of NMs

Due to its role as a RhoA GEF and its rosette center localized expression pattern, we hypothesized that *mcf2lb* may play a role in regulating rosette formation in the pLLP follower cells. To assess the role of *mcf2lb*, we generated two distinct *mcf2lb* mutant lines using CRISPR/Cas9-mediated gene editing (Supplemental Fig. 3A). Both lines are presumably loss-of-function and do not exhibit any significant phenotypic differences from each other (Supplemental Fig. 3B). Homozygous *mcf2lb* mutants are viable and fertile. As *mcf2lb* is maternally contributed (Supplemental Fig. 3C), we performed all our experiments in maternally zygotic homozygous mutants.

We next used Tg(-8.0*claudinB: lynGFP)^zf106^* transgene to assess the gross morphology of the pLL in the *mcf2lb* mutant embryos. At 3 dpf, *mcf2lb* mutant embryos showed an excess number of deposited trunk NMs compared to control embryos (Fig. 3A, B; average number of NMs in WT = 5.375 vs. *mcf2lb* mutants = 7.750; n = 8 WT embryos and n = 12 *mcf2lb* mutant embryos; p = 0.0002 by unpaired t-test). Due to the increase in the number of deposited NMs, we next asked whether the size of the NMs was different between WT and *mcf2lb* mutant embryos. To address this, we stained 3 dpf embryos with DAPI and used the Imaris cells feature to count the number of cells in each NM. Overall, *mcf2lb* mutants exhibited significantly smaller NMs than WT embryos (Fig. 3D - F; average size of NMs in WT = 36 cells vs. *mcf2lb* mutant = 20.65 cells; WT, n = 14 NMs from 4 embryos and *mcf2lb* mutant = 26 NMs from 5 embryos; p>0.0001 by unpaired t-test). Interestingly, no significant difference was observed in size of the first deposited NM when comparing WT to *mcf2lb* mutant embryos (Fig 3D, E, G; average size of first NM in WT = 37.50 cells vs. *mcf2lb* mutant = 25.80 cells; WT n = 4 NMs from 4 embryos and 5 NMs from 5 embryos; p = 0.11 by unpaired t-test). As the rostral-most pLLP rosette (first proto-NM) is patterned prior to the onset of migration (Nechiporuk and Raible, 2008), this indicates that *mcf2lb* does not play a role in regulating NM size until pLLP migration begins.

**Figure 3:**
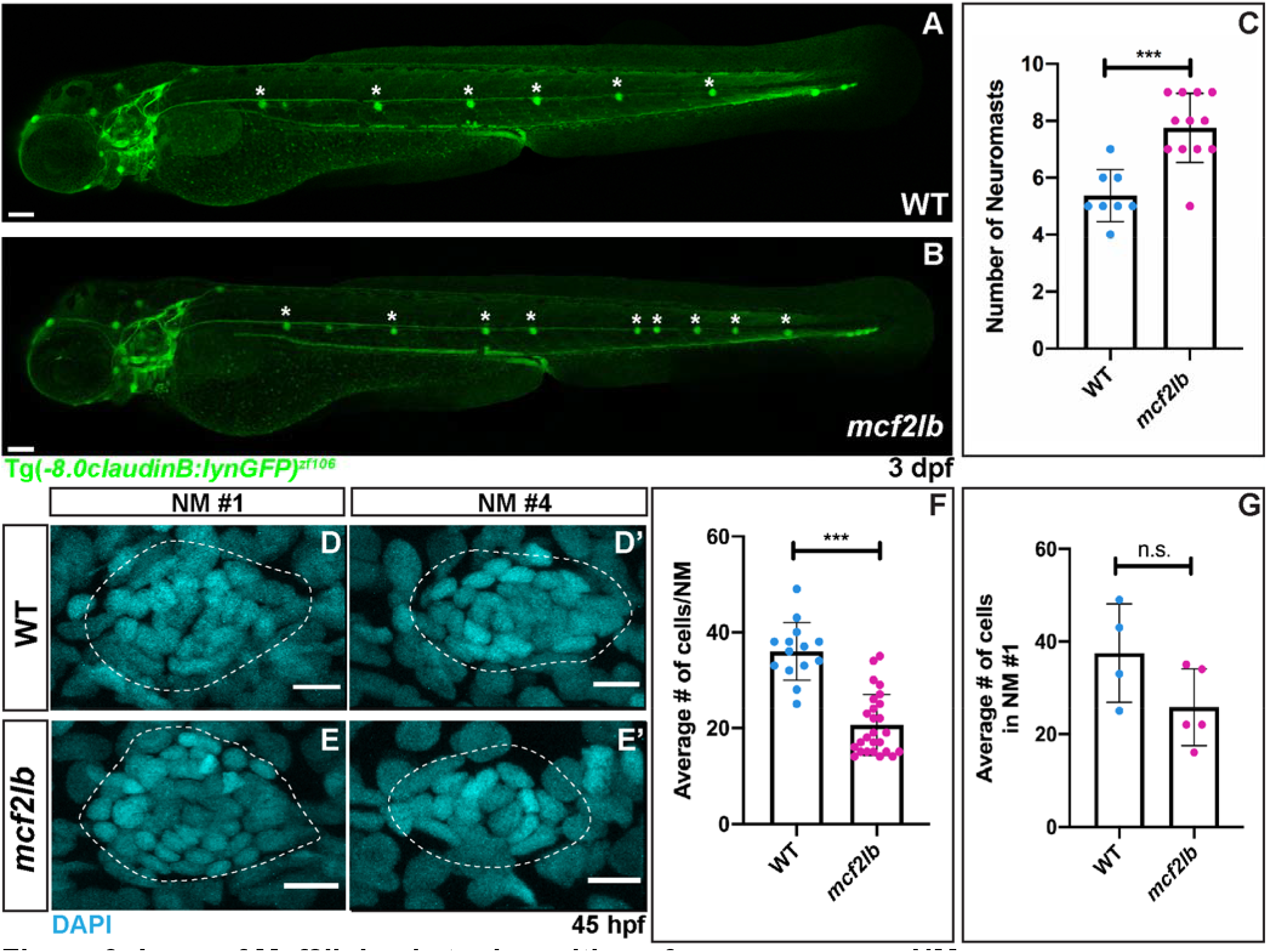
Loss of Mcf2lb leads to deposition of supernumerary NMs. (A, B) Tg(-8.0*claudinB: lynGFP)*^zf106^ transgene marks NMs in WT and *mcf2lb* mutant embryos 3 dpf. Note the excess number of deposited trunk NMs in the *mcf2lb* mutants. Asterisks mark deposited trunk NMs. (C) Average number of trunk NMs in WT (n = 8 embryos) and *mcf2lb* mutant (n = 12 embryos) embryos at 3 dpf. (D, E) DAPI staining of nuclei in deposited NMs in WT and *mcf2lb* mutants. (F) Average number of cells per NM in WT (n = 14 NMs from 4 embryos) and *mcf2lb* mutants (n = 26 NMs from 5 embryos). (G) Average number of cells in the first NM in WT (n = 4 NMs from 4 embryos) and *mcf2lb* mutants (n = 5 NMs from 5 embryos). *** – *p*<0.001 (unpaired t-test). Scale bars in A, B = 100 μm. Scale bars in D, E = 5 μm.

As there was an excess number of deposited NMs in *mcf2lb* mutants, we next asked whether hair cells are properly differentiated in *mcf2lb* mutant NMs. To address this question, we performed in situ hybridization to examine the expression of *atoh1a* and *deltaA*, two markers of hair cell precursors (Itoh and Chitnis, 2001). In WT, *atoh1a* and *deltaA* are expressed in one to two cells of opposing orientation in the deposited NMs (Supplemental Fig. 4). Similarly, *atoh1* and *deltaA* expression is limited to one or two cells in the deposited NMs in *mcf2lb* mutants (Supplemental Fig. 4). These results indicate that although there is an excess number of deposited NMs in *mcf2lb* mutants, NM hair cells still differentiate properly.

### *mcf2lb* mutants show impaired pLLP deposition behavior and abnormal pLLP organization

To address the cellular mechanism that leads to extra NMs, we next assayed NM deposition and pLLP organization in *mcf2lb* mutants. To achieve this, we imaged pLLP migration in WT or *mcf2lb* mutant Tg(-8.0*claudinB: lynGFP)^zf106^*- positive embryos between 30 and 44 hpf (Fig. 4A, B; Movie 1, 2). *mcf2lb* pLLP deposited multiple rosettes concurrently, instead of individual rosettes 5-7 somites apart as in WT embryos (Fig. 4B; Movie 1, 2). These groups of rosettes were often deposited as one large cluster of cells that then resolved into individual rosettes over time (Fig. 4B; Movie 2). Despite this abnormal behavior, velocity of the mutant pLLP did not differ than that of the WT (Fig. 4E; average velocity WT = 0.014 μm/s vs. *mcf2lb* mutant = 0.012 μm/s; n = 7 WT pLLP from 7 embryos and n = 6 *mcf2lb* mutant pLLP from 6 embryos; p = 0.45 by unpaired t-test). Additionally, there was no difference in the pLLP length (Fig. 4C; average length WT = 126.1 μm vs. *mcf2lb* mutant 131.8 μm; n = 7 WT pLLP from 7 embryos and n = 6 *mcf2lb* mutant pLLP from 6 embryos; p = 0.52 by unpaired t-test) or the number of cells within the migrating pLLP when comparing WT to *mcf2lb* mutants (Fig. 4D; average number of cells WT = 82.47 cells vs. *mcf2lb* mutant = 92.37 cells; n = 19 WT pLLP from 19 embryos and n = 19 *mcf2lb* mutant pLLP from 19 embryos; p = 0.14 by unpaired t-test). These results indicate that while the size of the pLLP and pLLP migration is not perturbed in *mcf2lb* mutants, rosette deposition behavior is impaired.

**Figure 4:**
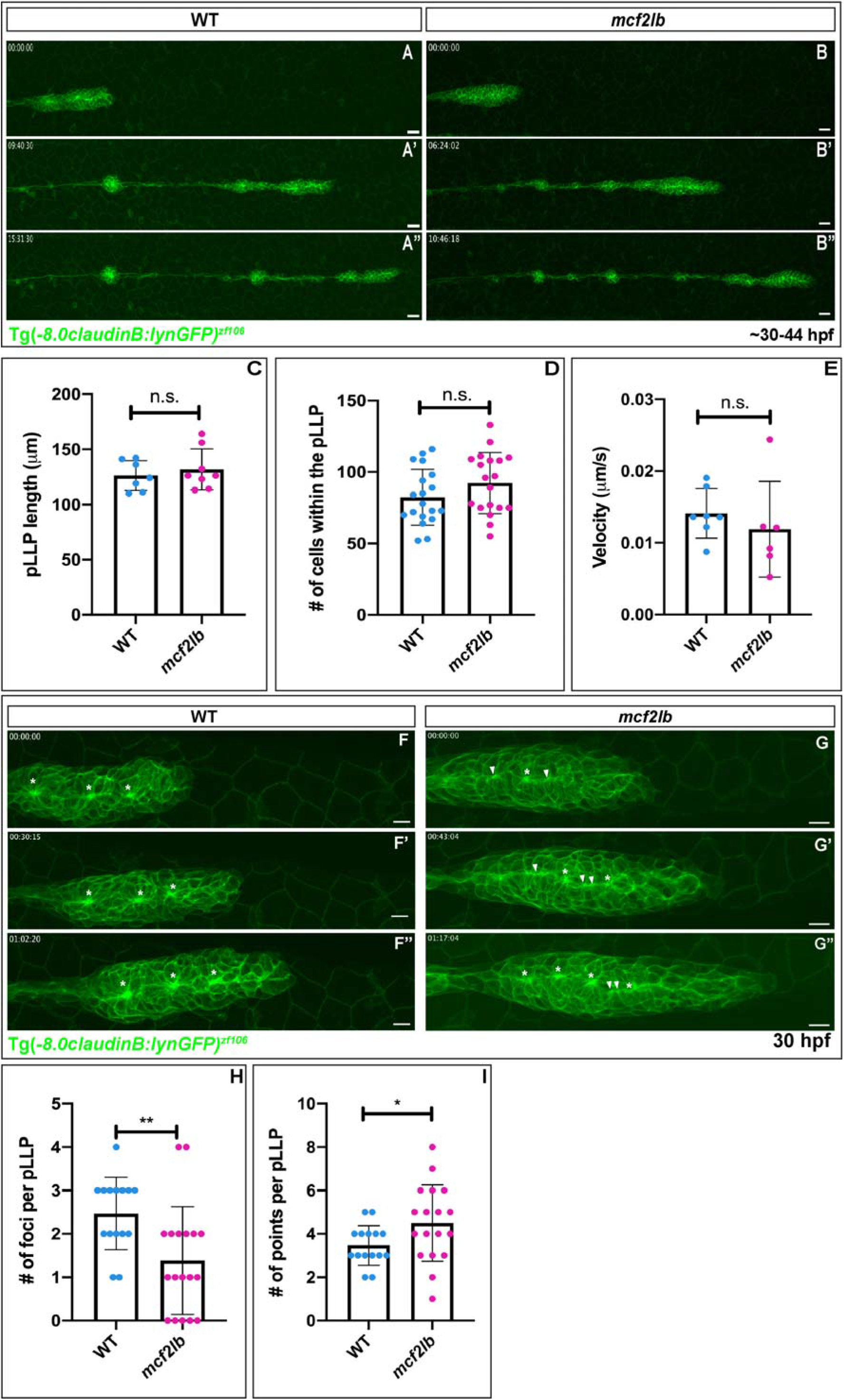
*mcf2lb* mutants show abnormal organization of the pLLP and NM deposition. (A, B) Lateral views of still images from time-lapse movies of either WT or *mcf2lb* mutant embryos positive for Tg(-8.0*claudinB: lynGFP)^zf106^* transgene (Movie 1, 2). Embryos were imaged from 30 to 44 hpf. (C) Average pLLP length in WT (n = 7 embryos) and *mcf2lb* mutants (n = 8 embryos). (D) Average number of cells within the pLLP in WT (n = 19 pLLP from 19 embryos) and *mcf2lb* mutants (n = 19 pLLP from 19 embryos). (E) Average velocity of the pLLP in WT (n = 7 pLLP from 7 embryos) and *mcf2lb* mutant (n = 6 pLLP from 6 embryos) embryos. (F, G) High-magnification lateral views of still images from movies of the migrating pLLP in WT and *mcf2lb* mutant embryos (Movie 3, 4). pLLPs were imaged for ∼1.5 hours starting ∼30 hpf. Note the abnormal organization of the pLLP in *mcf2lb* mutants and the beads on a string appearance along the midline instead of distinct clusters of membrane as in WT (Movie 3, 4). Asterisks = foci. Arrowheads = points. (H) Average number of foci per pLLP in WT (n = 15 pLLP from 15 embryos) and *mcf2lb* mutants (n = 18 pLLP from 18 embryos). Average number of points per pLLP in WT (n = 15 pLLP from 15 embryos) and *mcf2lb* mutants (n = 18 pLLP from 18 embryos) embryos. * – p<0.05, ** – p<0.01 (unpaired t-test). Scale bars in panels A, B = 20 μm. Scale bars in F, G, H = 10 μm.

Abnormal rosette deposition behavior implied that pLLP organization might be abnormal in *mcf2lb* mutants. To examine pLLP organization, we imaged pLLPs in either WT or *mcf2lb* mutant Tg(- 8.0*claudinB: lynGFP)^zf106^*-positive embryos at high magnification during migration. This revealed abnormal organization of rosettes in the pLLP of *mcf2lb* mutants (Fig. 4F, G; Movie 3, 4). WT pLLP usually contained 2-3 rosettes that display prominent gatherings of apical membrane, which we termed foci (Fig. 4F; Movie 3). In contrast, *mcf2lb* mutants displayed a “beads on a string” appearance of constricted membranes: the foci are not distinct and instead vary in size and shape (Fig. 4G; Movie 4). To quantify this pLLP phenotype, we divided the gatherings of membrane into two categories, either foci or points. Gatherings of membrane were deemed foci if width and the height of the focus were within one standard deviation above or below the average width and height of all foci measured in WT. If it did not meet those criteria, the gathering of membrane was deemed a point. In panels 4F and G, asterisks depict foci and arrowheads depict points. On average, *mcf2lb* mutants contained significantly fewer foci (Fig. 4H; average number of foci in WT = 2.47 foci vs. *mcf2lb* mutant = 1.39 foci; n = 15 WT pLLP from 15 embryos and n = 18 *mcf2lb* mutant pLLP from 18 embryos; p = 0.005 by unpaired t-test) and significantly greater number of points (Fig. 4I; average number of points WT = 3.47 vs. *mcf2lb* mutant = 4.50; n = 15 WT pLLP from 15 embryos and n = 18 *mcf2lb* mutant pLLP from 18 embryos; p = 0.049 by unpaired t-test) than WT pLLP. These results indicate that *mcf2lb* mutant pLLPs have abnormal cellular organization and a diminished ability to form proper rosettes.

### *mcf2lb* mutants show impaired apical constriction and rosette integrity in the pLLP

Given the observed rosette disorganization and enhanced expression of *mcf2lb* in rosettes, we hypothesized that apical constriction could be impaired in *mcf2lb* mutants. To examine this, we used Imaris to reconstruct surfaces of all cells in the pLLP in both WT and *mcf2lb* mutant pLLP (Fig. 5A, B). Following 3D cellular reconstruction, we examined the morphology of cells within the trailing region of the pLLP and found that many of the cells in the trailing region were not properly apically constricted in the *mcf2lb* mutants (Fig. 5C, D). Additionally, while in WT pLLP, cells usually constrict to an individual focus, we found that in *mcf2lb* mutants, some cells were making contacts with multiple points (Fig. 5E). To characterize these deficiencies, we divided cells into four categories: incorporated into rosettes, not incorporated into rosettes, touching multiple points, and those we could not include in analysis (dividing cells and cells that were not columnar). In *mcf2lb* mutants, the proportion of cells that was incorporated into rosettes was diminished in comparison to WT pLLP, whereas the percentage of cells that was not incorporated into rosettes was expanded (Fig. 5F; incorporated into rosettes: WT pLLP = 64% vs. *mcf2lb* mutant pLLP = 45%; p = 0.016 by unpaired t-test; not incorporated into rosettes: WT pLLP = 16% vs. *mcf2lb* mutant pLLP = 30%; n = 4 WT pLLP from 4 embryos and n = 4 *mcf2lb* mutant pLLP from 4 embryos; p = 0.03 by unpaired t-test). Additionally, the percentage of cells touching multiple points was expanded in *mcf2lb* mutant pLLP in comparison to WT pLLP (Fig. 5F; WT pLLP = 3% vs. *mcf2lb* mutant = 10%; n = 4 WT pLLP from 4 embryos and n = 4 *mcf2lb* mutant pLLP from 4 embryos; p = 0.05 by unpaired t-test). To quantify apical constriction, we measured the apical constriction index (ACI) of cells incorporated into rosettes in the trailing region of both WT and *mcf2lb* mutant pLLPs. In *mcf2lb* mutants, the average ACI of cells incorporated into rosettes was significantly increased in comparison to WT pLLP (Fig. 5G; WT = 0.50 vs. *mcf2lb* = 0.65; WT n = 169 cells from 4 pLLP from 4 embryos and *mcf2lb* = 139 cells from 4 pLLP from 4 embryos; p<0.0001 Mann-Whitney U). To further examine this impairment, we also compared the average apical width and basal width of cells incorporated into rosettes. We found a significant increase in apical width in *mcf2lb* mutant cells (Fig. 5H; WT = 2.28 μm vs. *mc2lb* mutants = 3.18 μm; WT n = 169 cells from 4 pLLP from 4 embryos *mcf2lb* = 139 cells from 4 pLLP from 4 embryos; p<0.0001 Mann-Whitney U). However, there was no significant difference in the basal width of cells when comparing WT to *mcf2lb* mutants (Fig. 5I; WT = 6.06 μm vs. *mc2lb* mutants = 5.94 μm; WT n = 169 cells from 4 pLLP from 4 embryos, *mcf2lb* mutant = 139 cells from 4 pLLP from 4 embryos; p = 0.55 Mann-Whitney U). These results indicate that in *mcf2lb* mutants, there is an impairment in the ability of cells incorporated into rosettes to apically constrict, while the basal size of cells is unchanged.

**Figure 5:**
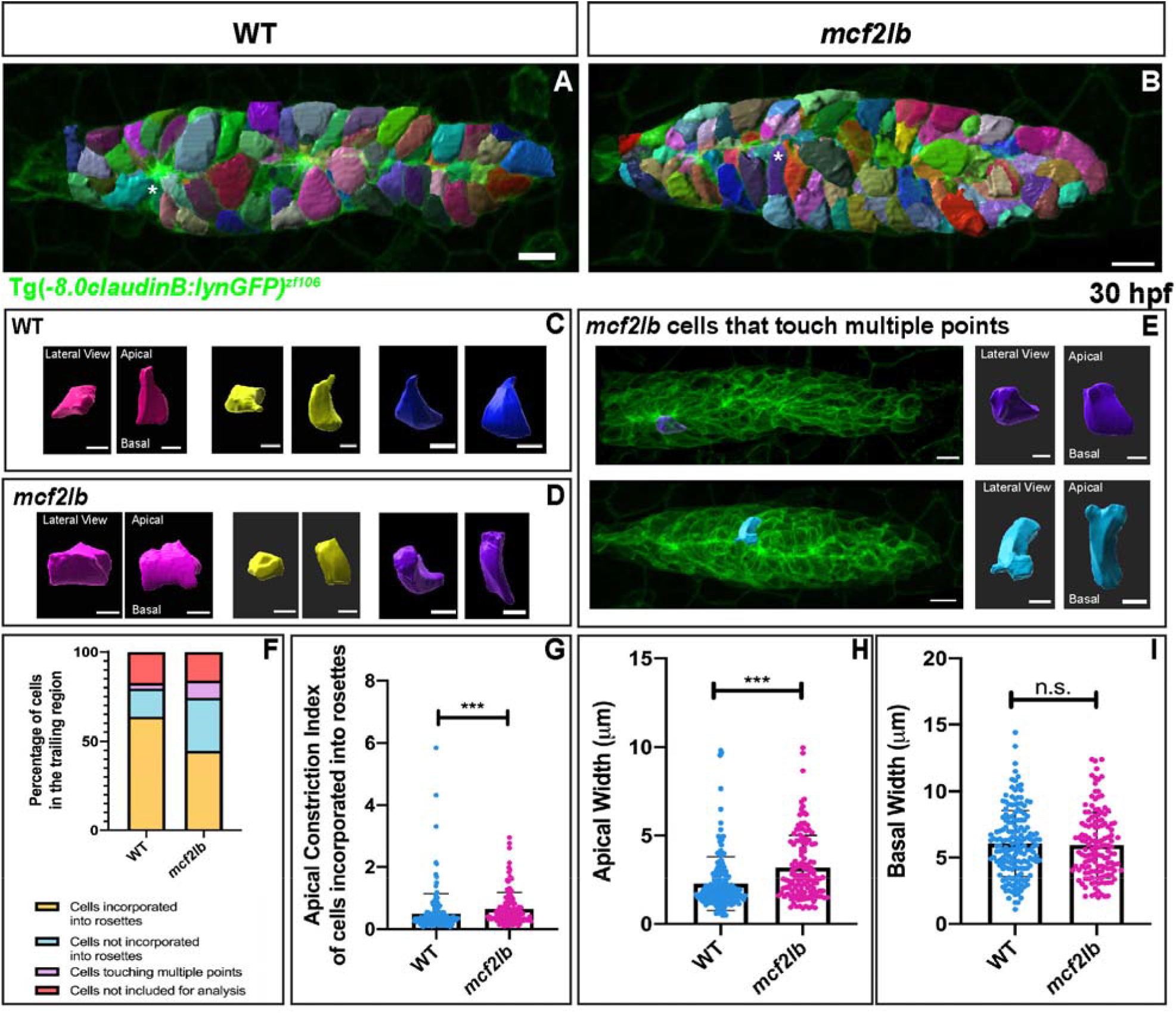
*mcf2lb* mutants show impaired apical constriction of cells incorporated into rosettes. (A, B) Cellular reconstruction of WT and *mcf2lb* mutant pLLPs. (C, D) Examples of apically constricted cells from WT (C) and *mc2lb* mutant (D) pLLPs. The left side panels show lateral (top-down) view as in panels A and B, and the right panels show apical/basal views of the cell (cell is virtually turned by 90 degrees). Blue and purple cells marked by the asterisks in panel A and B are also shown in C and D, respectively. (E) Examples of cells that are making contact with multiple points in *mcf2lb* mutant embryos. Left panel is a lateral (top-down) view; right panel is apical/basal view. (F) Categorical breakdown of cells in the trailing region of WT and *mcf2lb* mutant pLLPs. Note that in *mcf2lb* mutants, there is an increase in the percentage of cells that touch multiple points and an increase in the percentage of cells not incorporated into rosettes. Subsequently, there is a decrease in the percentage of cells that are incorporated into rosettes. (G) Average apical constriction index of cells that are incorporated into rosettes in WT (n = 169 cells from 4 pLLP from 4 embryos) and *mcf2lb* mutant (n = 139 cells from 4 pLLP from 4 embryos) pLLPs. (H) Average apical width of cells incorporated into rosettes in WT (n = 169 cells from 4 pLLP from 4 embryos) and *mcf2lb* mutant (n = 139 cells from 4 pLLP from 4 embryos) pLLPs. (I) Average basal width of cells incorporated into rosettes in WT (n = 169 cells from 4 pLLP from 4 embryos) and *mcf2lb* mutant (n = 139 cells from 4 pLLP from 4 embryos) pLLPs. Note the apical width is significantly increased in *mcf2lb* mutants whereas there is no significant difference in the basal width. *** – p<0.0001 (Mann – Whitney U test) Scale bars for panels A, B = 10 μm; panels C, D = 5 μm; panel E pLLP = 10 μm, individual cells = 5 μm.

Previous studies revealed that the rostral most rosette in the pLLP forms a microlumen within its apical region (Durdu et al., 2014). Secreted Fgf ligands accumulate in the microlumen and maintain rosette integrity and orderly NM deposition (Durdu et al., 2014). After rosette deposition, the microlumen eventually expands into a lumen housing hair cells’ stereo- and kono-cilia. To assess, whether microlumen formation is affected in *mcf2lb* mutants, we transiently expressed secreted GFP (secGFP) in the migrating pLLP as well as newly deposited NMs. In wildtype controls, even a single cell with secGFP expression was sufficient to mark the microlumen in the trailing rosettes and recently deposited NMs (Supplemental Fig. 5A, C). In contrast, we detected low secGFP in *mcf2lb* mutants (Supplemental Fig. 5A – E; average normalized fluorescence intensity in WT = 0.08433 vs. *mcf2lb* mutant = 0.01883; n = 7 WT and n = 6 *mcf2lb;* p = 0.0003 by unpaired t-test). These data indicate that the trailing rosette and, consequently, NM integrity is compromised in the *mcf2lb* mutant.

### *mcf2lb* mutants display abnormal apical membrane dynamics

In order to visualize the dynamics of apical membranes in single pLLP cells, we generated mosaic primordia that contained a few cells with fluorescently labeled membranes. To achieve this, we transplanted a small number of cells from a donor embryo expressing the Tg(*prim:lyn2-mCherry*) transgene into a Tg(-8.0*claudinB: lynGFP)^zf106^* host. Observation of WT cells in a WT background revealed that once incorporated into a rosette, apically constricted cells maintain contact with the rosette center during migration (Fig. 6A; Movie 5). Interestingly, we observed some apical membrane expansion and retraction in apically constricted cells both in WT and mutant pLLPs. To quantify apical membrane dynamics, we measured “average membrane variability” and average apical membrane width (Fig. 6C, D). We defined “average membrane variability” as the standard deviation of the apical width of a cell over the course of an hour imaging period. *mcf2lb* mutant cells transplanted into the mutant background revealed that the apical membranes of mutant cells are much more variable than WT cells over the course of the imaging period (Fig. 6B – D; Movie 6). When quantified, the average membrane variability in WT cells was 1.19 μm (n = 22 cells from 7 embryos) vs. 2.01 μm in *mcf2lb* mutant cells (n = 13 from 3 embryos) (p = 0.01 Mann-Whitney U). These results indicate that *mcf2lb* mutant cells cannot maintain an apically constricted state.

**Figure 6:**
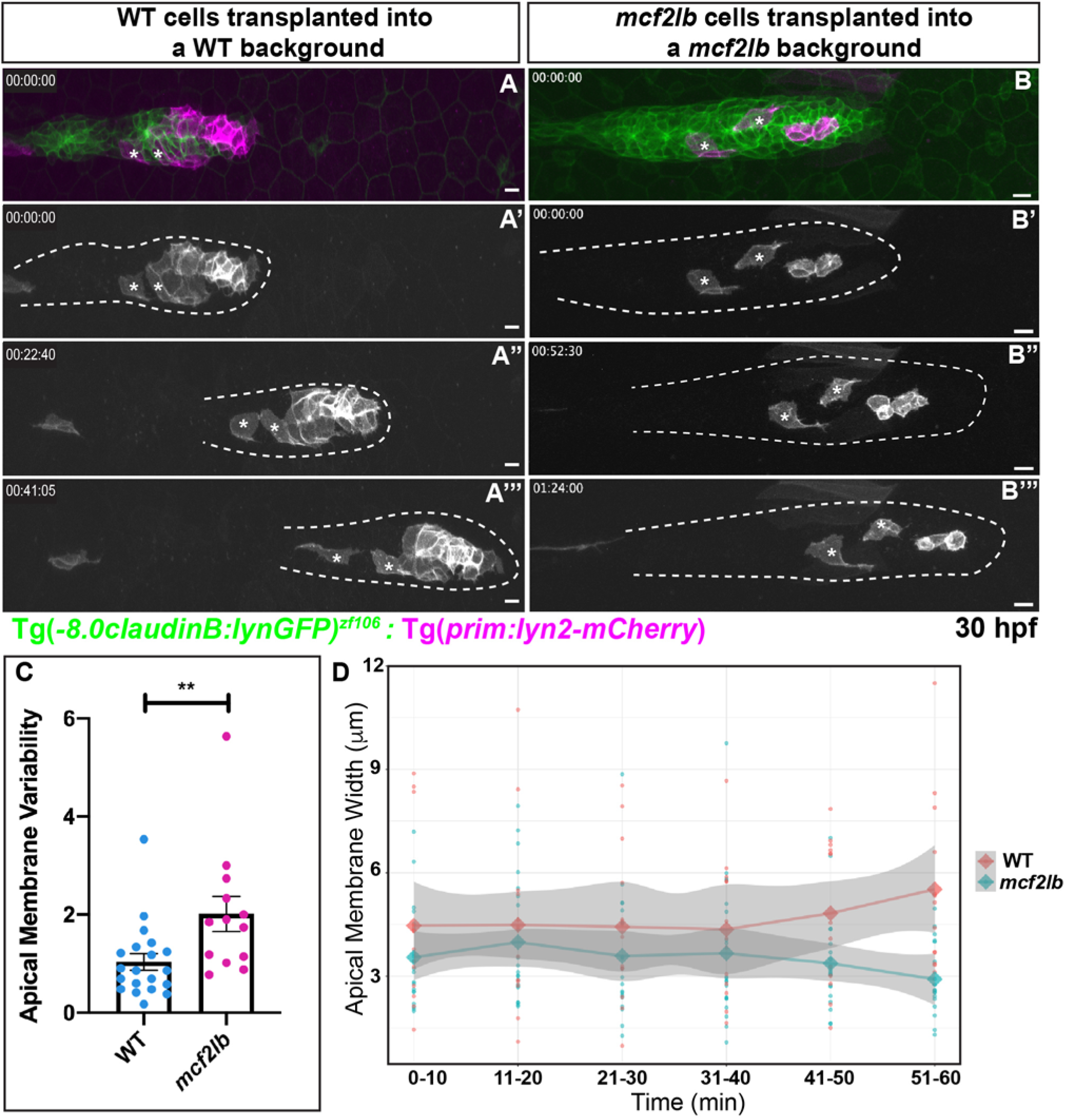
*mcf2lb* mutants show greater variability in apical membrane dynamics. (A, B) Donor cells derived from either WT or *mcf2lb* mutant embryos Tg (*prim:lyn2-mCherry*; magenta) were transplanted into WT or *mcf2lb* mutant Tg(-8.0*claudinB: lynGFP;* green)^zf106^ positive embryos, respectively, and mounted for live imaging between 30 and 32 hpf (Movie 5, 6). Tg(*prim:lyn2-mCherry*) cells are shown in gray scale for clarity. Asterisks indicate cells used for analysis (C) Average membrane variability of transplanted WT (n = 22 cells from 7 embryos) or *mcf2lb* mutant (n = 13 cells from 3 embryos) cells over time from movies obtained from experiments in panels A and B. Membrane variability is defined as the standard deviation of the apical width of a cell over an hour period of imaging in apically constricted cells. (D) Apical membrane width over time with SEM. Note higher variability in the mutant. ** – p<0.01 (Mann-Whitney U test). Scale bars for panels A, B = 10 μm.

### Cell polarity is largely normal in *mcf2lb* mutant pLLP

As *mcf2lb* mutant cells show impairment in apical constriction, we next asked whether mutant pLLPs were properly polarized. To examine this, we visualized localization of the tight junction scaffolding protein ZO-1 in WT and *mcf2lb* mutant pLLPs at 45 hpf (Niessen, 2007). In WT pLLP, ZO-1 staining showed enhancement at the rosette centers (Fig. 7A). Additionally, in the caudal most rosette, ZO-1 formed a ring-like structure (Fig. 7A). After digitally rotating the pLLP 90 degrees, ZO-1 immunostaining appeared apically localized at the rosette centers (Fig. 7A’). In *mcf2lb* mutant pLLP, ZO-1 remained localized to the midline, however, there was no ring-like structure present in the most distal rosette (Fig. 7F). Instead, ZO-1 staining was more punctate (Fig. 7F). Nevertheless, ZO-1 remained apically localized in *mcf2lb* mutants (Fig. 7F’ K; percentage of ZO-1 signal apically localized WT = 53% vs. *mcf2lb* mutant = 44%; WT n = 8 pLLP from 8 embryos, *mcf2lb* mutant n 13 pLLP from 13 embryos; p = 0.17 by unpaired t-test). These results indicate that while the organization of ZO-1 appears to be slightly impaired, cells in the pLLP in *mcf2lb* mutants still are properly polarized.

**Figure 7:**
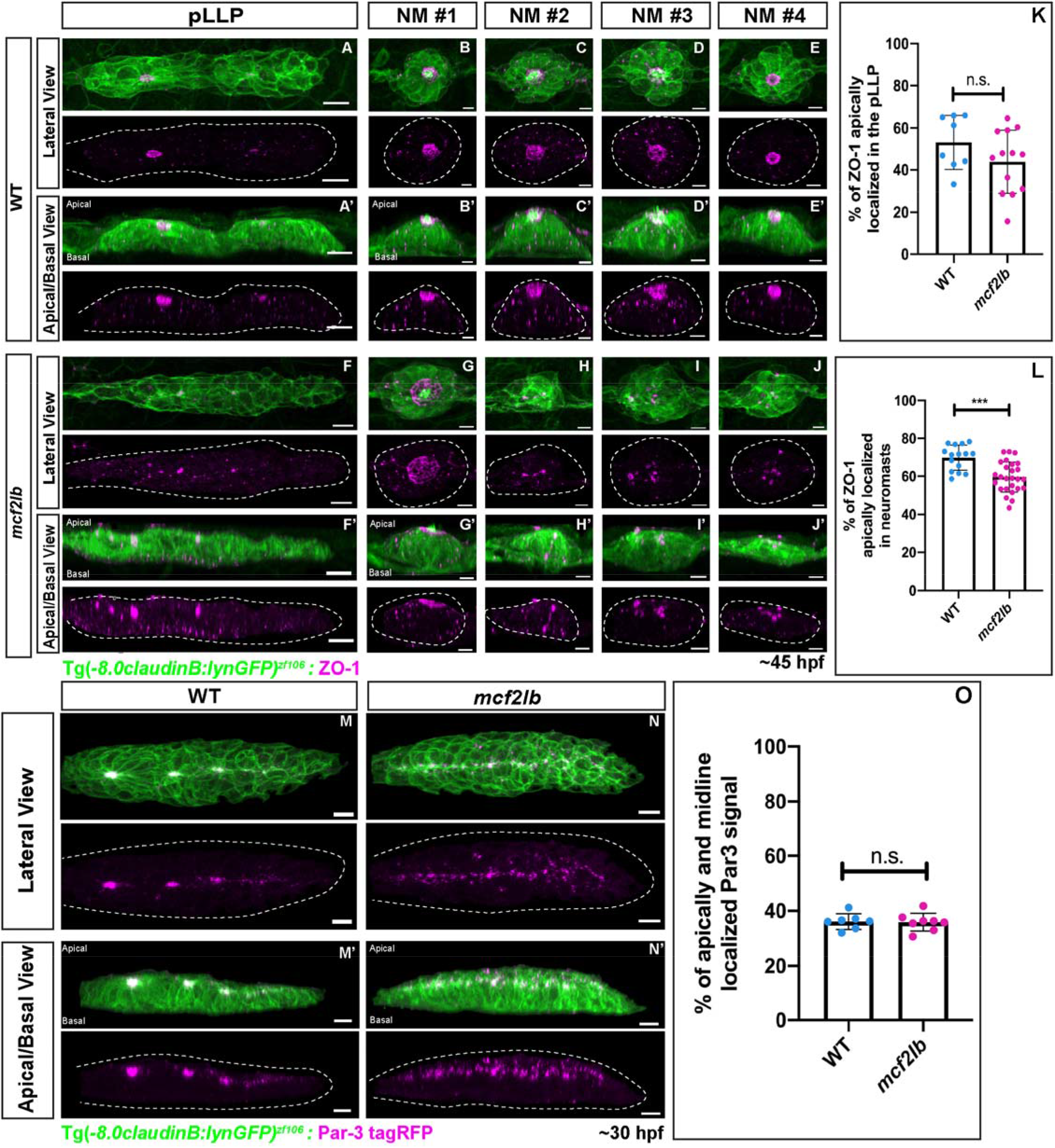
pLLP cell polarity is largely unaffected in *mcf2lb* mutants. (A – J) Immunostaining for the tight junction marker ZO-1 in WT and mutant pLLPs at 45 hpf. Top panels show lateral (top-down) view, bottom panels show an apical-basal view (images were virtually rotated 90 degrees). Dotted lines indicate pLLP in B and F and NMs in B – E and G – J. Note the ring structure in the trailing-most rosette and the apical localization of ZO-1 signal in WT. Similarly, the ring structure and the apical localization of ZO-1 signal is present in all deposited NMs. In contrast, ZO-1 signal in the pLLP and within the NMs is disorganized in *mc2lb* mutants. (K) Quantification of the percentage of apically localized ZO-1 signal in the pLLP in WT (n = 8 pLLP from 8 embryos) and *mcf2lb* mutant (n = 13 pLLP from 13 embryos) embryos. (L) Quantification of the percentage of apically localized ZO-1 signal in neuromasts in WT (n = 15 NMs from 4 embryos) and *mc2lb* mutant (n = 26 NMs from 5 embryos) embryos. (M, N) Par-3-tagRFP expression in WT and *mcf2lb* mutant pLLPs. Note Par-3 localization to the rosette centers and the midline in WT embryos; in contrast, Par-3 is localized to the midline but not organized into rosette centers in *mcf2lb* mutants. Par-3 is apically localized in both WT and *mcf2lb* mutant pLLPs. (O) Quantification of the percentage of apically- and midline-localized Par-3 signal in WT (n = 7 pLLP from 7 embryos) and *mcf2lb* mutant (n = 8 pLLP from 8 embryos) pLLPs. *** – p<0.001 (unpaired t-test). Scale bars for panels A, F, M, N = 10 μm and panels B – D, G – J = 5 μm.

In addition to ZO-1, we also examined the localization of the polarity marker Par-3. Par-3 is a component of the aPKC complex that assembles apically to tight junctions in epithelial cells (Chen and Macara, 2005). To visualize Par-3’s localization, we injected *par3-tagRFP* RNA into either WT or *mcf2lb* mutant Tg(-8.0*claudinB: lynGFP)^zf106^*- positive embryos. Embryos were mounted at 30 hpf for live imaging to examine Par-3-tagRFP localization. In WT, Par-3 localized to the rosette centers in the migrating pLLP (Fig. 7M). Digital rotation of the images by 90 degrees revealed Par-3 apical localization in WT pLLP (Fig. 7M’). In *mcf2lb* mutant pLLP, Par-3 localized entirely along the pLLP midline instead of rosette centers (Fig. 7N). Digital rotation of the images 90 degrees revealed proper apical localization of Par-3 (Fig. 7N’ O; percentage of Par-3 signal that is midline and apically localized, WT = 36% vs. *mcf2lb* mutant = 36%; WT n = 7 pLLP from 7 embryos, *mcf2lb* mutant n = 8 pLLP from 8 embryos; p = 0.88 by unpaired t-test). These results, together with the ZO-1 localization, indicate that cells within *mcf2lb* mutant pLLP are normally polarized, but show impairment in their organization into rosettes.

Finally, we examined ZO-1 immunostaining in deposited NMs at 45 hpf in WT and *mcf2lb* mutant embryos. As NMs mature, a ring-like opening forms at the center of a maturing NM to allow for stereociliary bundles to push through (Fig. 7B’). In WT embryos, all deposited NMs had a ring of ZO-1 that was apically localized (Fig. 7B – E). In *mcf2lb* mutants, this ring structure was only observed in NM #1 (Fig. 7G). The remaining NMs showed disorganized ZO-1 staining and impaired apical localization of ZO-1 (Fig. 7H-J,L; percentage of signal apically localized WT = 69% vs. *mcf2lb* mutant = 60%; WT n = 15 NMs from 4 embryos, *mcf2lb* mutant n = 26 NMs from 5 embryos; p = 0.0002 by unpaired t-test). These results indicate that abnormal apical constriction in mutant rosettes ultimately leads to the formation of disorganized NMs.

### RhoA signaling is disrupted in *mcf2lb* mutant pLLP

Mcf2l has been shown to act as RhoA GEF in *in vitro* contexts in mammalian cells and cortical neurons (Whitehead et al., 1999; Hayashi et al., 2013). Additionally, RhoA signaling components necessary for proper apical constriction, Rock and non-muscle Myosin II, have been shown to be activated downstream of Mcf2l (Liu et al., 2006). Since the RhoA pathway is required for apical constriction of pLLP cells (Harding and Nechiporuk, 2012), we next asked whether downstream effectors of RhoA signaling are disrupted in *mcf2lb* mutant pLLP. In the pLLP, RhoA activates the Rho kinase Rock-2a, which is apically scaffolded by Schroom3 (Ernst et al., 2012; Harding and Nechiporuk, 2012). Rock-2a then phosphorylates Myosin Regulatory Light Chain (MRLC), which activates non-muscle Myosin-II-mediated apical constriction (Harding and Nechiporuk, 2012). We first examined whether Rock-2a is present and properly localized. Immunostaining of Rock-2a in WT pLLP showed accumulations of Rock-2a at rosette centers (Fig. 8A). The overall levels of Rock-2a signal were not significantly different when comparing overall fluorescence per cell in WT and *mcf2lb* mutant pLLPs. However, there was a significant difference in the fluorescence intensity at the rosette centers when comparing WT to *mcf2lb* mutants (Fig. 8A-D; average fluorescent intensity of Rock-2a per cell in WT pLLP = 4185000 vs. *mcf2lb* mutant pLLP = 5080000; WT n = 10 pLLP from 10 embryos and *mcf2lb* mutant n = 10 pLLP from 10 embryos; p = 0.19 by unpaired t-test; average fluorescent intensity of Rock-2a at rosette centers WT = 1198500 vs. *mcf2lb* mutant = 931700; WT n = 16 rosette centers from 10 pLLP from 10 embryos and *mcf2lb* mutant n = 30 rosette centers from 10 pLLP from 10 embryos; p = 0.003 by unpaired t-test). To examine apical localization of Rock-2a, images were digitally rotated 90 degrees. In WT and *mcf2lb* mutant pLLPs, Rock-2a was localized apically (Fig. 8A’,B’). These results demonstrate that while Rock-2a levels and its apical localization are not changed, there is a decreased amount of Rock2a at the rosette centers in *mcf2lb* mutants.

**Figure 8:**
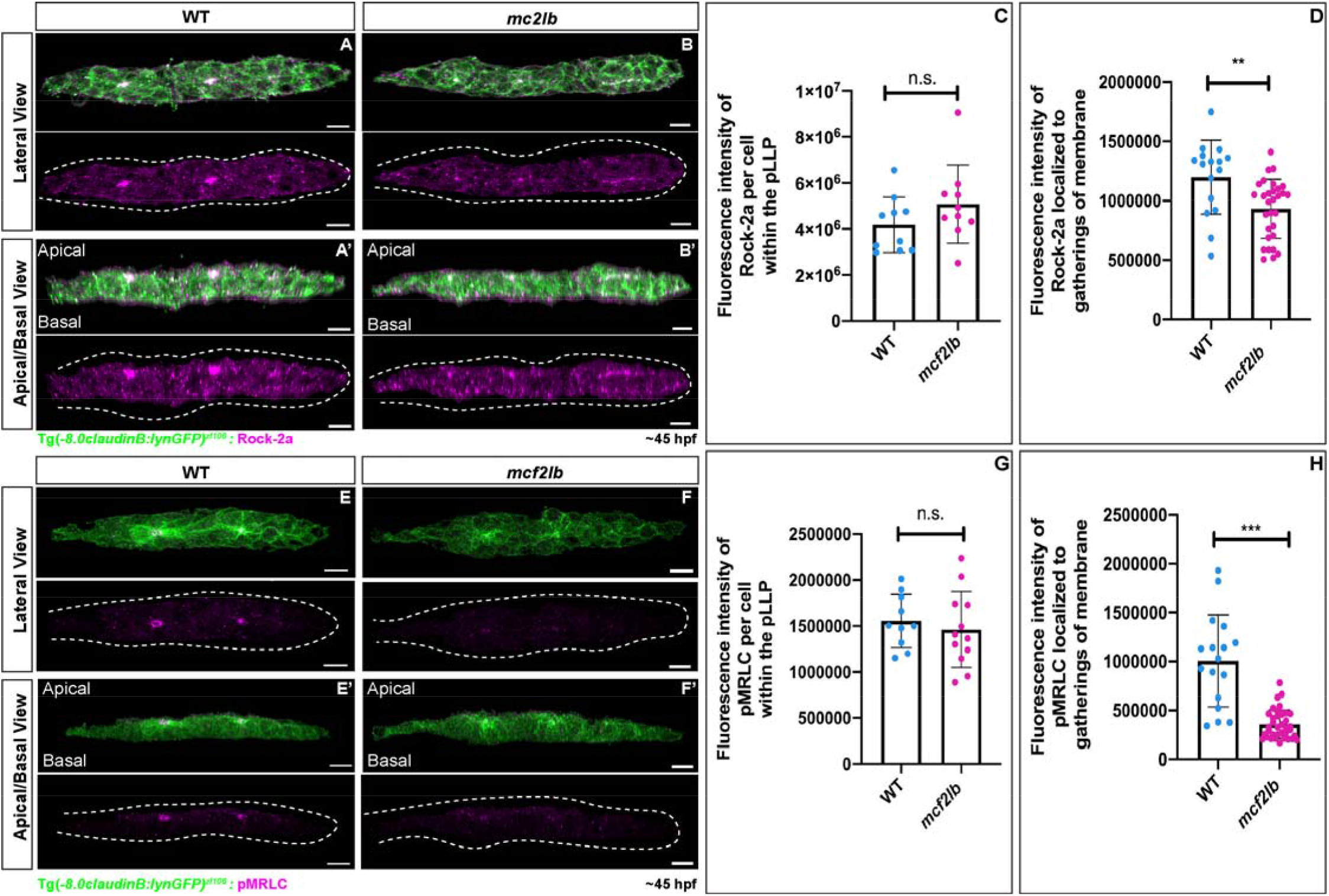
RhoA signaling is disrupted in *mcf2lb* mutants. (A, B) Immunostaining for Rock-2a in WT and *mcf2lb* mutants. Top panels show lateral view (top-down), bottom panels show apical-basal view (images were virtually rotated 90 degrees). Images are masked to only show signal within the pLLP. Dotted lines indicate pLLP. Note the localization of Rock-2a is diminished in *mcf2lb* mutant pLLP. (C) Total fluorescence intensity of Rock-2a per cell in WT (n = 10 pLLP from 10 embryos) and *mcf2lb* mutant (n = 10 pLLP from 10 embryos) pLLPs. (D) Fluorescence intensity of Rock-2a at rosette centers in WT (n = 16 points from 10 pLLP from 10 embryos) and *mcf2lb* mutant (n = 30 points from 10 pLLP from 10 embryos) pLLPs. (E, F) Immunostaining for pMRLC in WT and *mcf2lb* mutant embryos. Top panels show lateral (top-down) views, bottom panels show apical-basal view. Images are masked to only show signal within the pLLP. Dotted lines indicate pLLP. Note diminished localization of pMRLC in the *mcf2lb* mutant pLLP. (G) Total fluorescence intensity of pMRLC per cell in WT (n = 10 pLLP from 10 embryos) and *mcf2lb* mutant (n = 13 pLLP from 13 embryos) pLLPs. (H) Fluorescence intensity of pMRLC at the gatherings of the membranes in WT (n = 17 points from 10 pLLP from 10 embryos) and *mcf2lb* mutant (n = 37 points from 13 pLLP from 13 embryos) pLLPs. ** – p<0.01 *** – p<0.001 (unpaired t-test). Scale bars in A, B, D, E = 10μm.

Because anti-Rock2a does not distinguish between active and inactive forms of the protein, we asked whether a downstream effector of Rock2a, the MRLC component of non-muscle Myosin II, is activated in the mutant pLLP. In WT pLLP, pMRLC accumulated at rosette centers (Fig 8E). However, in *mcf2lb* mutants, pMRLC signal was diminished at rosette centers (Fig. 8F, H; WT = 1005529 vs. *mcf2lb* = 357595; WT n = 17 rosette centers from 10 pLLP from 10 embryos, *mcf2lb* mutant = 37 rosette centers from 10 pLLP from 10 embryos; p<0.0001 by unpaired t-test). To examine apical localization of pMRLC, images were digitally rotated 90 degrees. In both WT and *mcf2lb* mutant pLLPs, pMRLC was properly apically localized (Fig. 8E’,F’). These results indicate that there is a significant reduction in the amount of active MRLC at the rosette centers, which ultimately leads to impaired apical constriction.

## DISCUSSION

Using scRNA-seq, we defined a comprehensive set of genes that regulate the actin cytoskeleton during pLLP migration. We then focused on *mcf2lb* and showed that it is required for apical constriction and rosette integrity during pLLP migration. As evidenced by reduced RhoA downstream effectors, Rock-2a and pMRLC, our study supports the role of Mc2lb as GEF for RhoA. We propose that Mc2lb activates RhoA, which subsequently activates Rock-2a; Rock-2a then phosphorylates non-muscle Myosin-II and induces apical constriction through the interaction of non-muscle Myosin-II with apically localized actin fibers.

### Follower cells have higher levels of actin regulatory genes

In silico analysis of genes that are involved in actin binding, Rho GTPases, Rho-GEFs, Rho-GAPs, and actin polymerization revealed higher expression in follower cells. This was particularly pronounced for actin polymerization, as all genes involved in this process showed higher expression in followers. This observation is consistent with a recent study that assayed stress forces along the migrating pLLP. It determined that the pLLP exerts higher stresses in the trailing rather than leading region (Yamaguchi et al., 2022). The authors hypothesize that the trailing end must therefore be “pushing” the pLLP instead of the leading cells “pulling” the pLLP forward (Yamaguchi et al., 2022). Future studies will examine the role of individual actin regulatory genes within the trailing region and how they potentially promote the “pushing” of the pLLP forward to facilitate pLLP migration.

### Improper apical constriction results in impaired rosette integrity and rosette deposition

Normal deposition of rosettes in the pLLP is necessary for the even spacing of pLL mechanosensory organs along the trunk. In *mcf2lb* mutants, we observe impaired rosette deposition behavior, with the deposition of large clusters of cells, that ultimately resolves into 2-3 NMs. While the mechanisms by which rosettes are deposited is not fully understood, two previous studies have identified some of the molecular mechanisms that facilitate deposition events (Aman et al., 2011; Durdu et al., 2014). Aman et al., (2011) showed that deposition events are not dependent on external cues such as somite boundaries (Aman et al., 2011), providing evidence that NM deposition is cell autonomous to the pLLP and not induced by external factors. The second study determined that proliferation during pLLP migration and subsequent lengthening of the pLLP is necessary for proper NM deposition (Aman et al., 2011). Disruption of cell proliferation through the slowing of the cell cycle resulted in NMs that were deposited farther apart despite pLLP migratory speed being maintained (Aman et al., 2011). These results suggest that size and length of the pLLP might be potential mediators of pLLP deposition behavior. Our study also showed that in addition to these two factors, rosette integrity is also required for normal NM deposition.

In addition, Fgf signaling is necessary for proper NM deposition (Durdu et al., 2014). Reducing Fgf signaling through the use of the Fgf inhibitor, SU5402, resulted in a dose-dependent delay in NM deposition, whereas increasing Fgf activity through overexpression of the Fgf ligand, Fgf3, resulted in an increased rate of NM deposition (Durdu et al., 2014). Although, in contrast to the *mcf2lb* mutant phenotype, this higher Fgf level did not result in more trunk NMs compared to controls. This led to the hypothesis where Fgf activity within the trailing pLLP can control the frequency of NMs deposition. As Fgf ligands are concentrated in the microlumen of the trailing most rosette (Durdu et al., 2014), disruption of this microlumen will affect the deposition of NMs (Durdu et al., 2014). Our experiments with secGFP implied that the microlumen is indeed disrupted in *mcf2lb* mutants, and, consistent with the above study, this ultimately leads to abnormal NM deposition.

### Is *mcf2lb* involved in the formation or the maintenance of rosettes in the pLLP?

Our study revealed that Mcf2lb regulates apical constriction and rosette integrity. Can we distinguish between whether Mcf2lb is involved in the formation and/or maintenance of rosettes? We would argue for the latter for the following reasons. We observe at least some cells that are properly apically constricted within mutant pLLP. In addition, our data showed that even in WT cells, the width of apical region in constricted cells fluctuates. This implies that this region is under tension and there is an active mechanism in place to maintain apical constriction. We believe this mechanism is, at least in part, mediated by Mcf2lb, because we observe a much wider range of apical region fluctuation in mutant cells compare to WT cells. Notably, mutant pLLP cells are still able to apically constrict to a certain extent and form rosettes of various sizes. This indicates that there are additional GEFs that regulate RhoA signaling to ensure proper apical constriction within the migrating pLLP. In summary, our data argues for Mcf2lb’s role in rosette maintenance, which is ultimately necessary for the formation of a functional sensory organ. In addition, our scRNA-seq data set provides a wealth of information to look for additional GEFs and GAPs that regulate RhoA-mediated apical constriction in the pLLP.

### Other roles for RhoA signaling in the pLLP

RhoA has numerous other reported roles in regulating the cytoskeleton outside its role in apical constriction. In certain contexts, RhoA is necessary for cellular adhesion, cell survival, cell division, and cell migration, and in particular regulating aspects of protrusion dynamics (Jaffe and Hall, 2005). Is it possible that in addition to its role in regulating RhoA during apical constriction, Mcf2lb may be mediating other cellular processes necessary for pLLP differentiation and /or migration? We found that migration speed was unchanged in *mcf2lb* mutants, implying that protrusive behavior is largely normal. We also showed that pLLP cells are polarized and we did not observe any phenotypes previously associated with the loss of cellular adhesion (Colak-Champollion et al., 2019; Matsuda and Chitnis, 2010). Finally, mutant pLLPs have the same cell number when compared with WT pLLPs, arguing that proliferation and survival is normal as well. Altogether, this indicates that in the pLLP, Mcf2lb is specifically involved in maintaining apical constriction rather than regulating other cellular processes.

## AUTHOR CONTRIBUTIONS

Conceived and designed experiments: HO, AM, AVN. Performed scRNA-seq experiments: AVN. Performed remaining experiments: HO, AM. Analyzed data for scRNA-seq: AVN, NC, LH. Analyzed remaining data: HO, AM. Wrote Manuscript: HO. Edited Manuscript: AVN, AM.

## ACKNOWLEDGEMENTS

The authors thank Dr. Holger Knaut and Dr. Darren Gilmour for reagents. This work was supported with funding provided to HO from the NICHD (F31HD095606; http://www.nichd.nih.gov) and to AVN from the NIGMS (R01GM130868); http://www.ninds.nih.gov).

## Notes

### Competing Interest Statement

The authors have declared no competing interest.

### Summary of Updates

An updated version of Figure 2.

