## Supplemental Figures for "RhoA GEF Mcf2lb regulates rosette integrity during collective cell migration"

**Supplemental Materials**

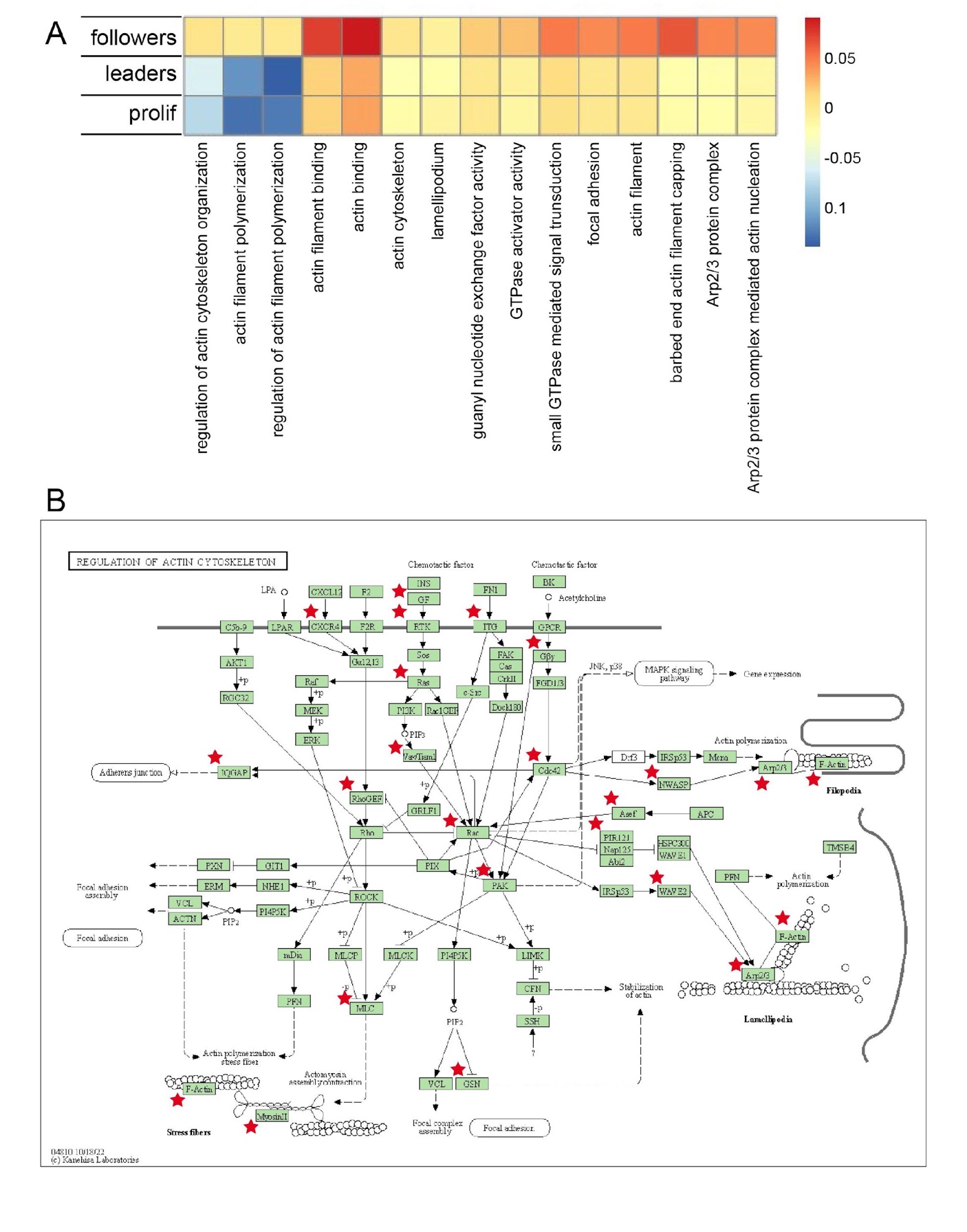

**Supplemental Figure 1: GO term and KEGG analysis of the scRNA-seq data set for genes that regulate actin dynamics.**

(A) Heatmap illustrating pLLP module scores for GO terms that are associated with various processes involving actin. (B) Regulation of actin cytoskeleton KEGG pathway map shows presence (red star) of molecules that are associated with of this process in our scRNA-seq data set.

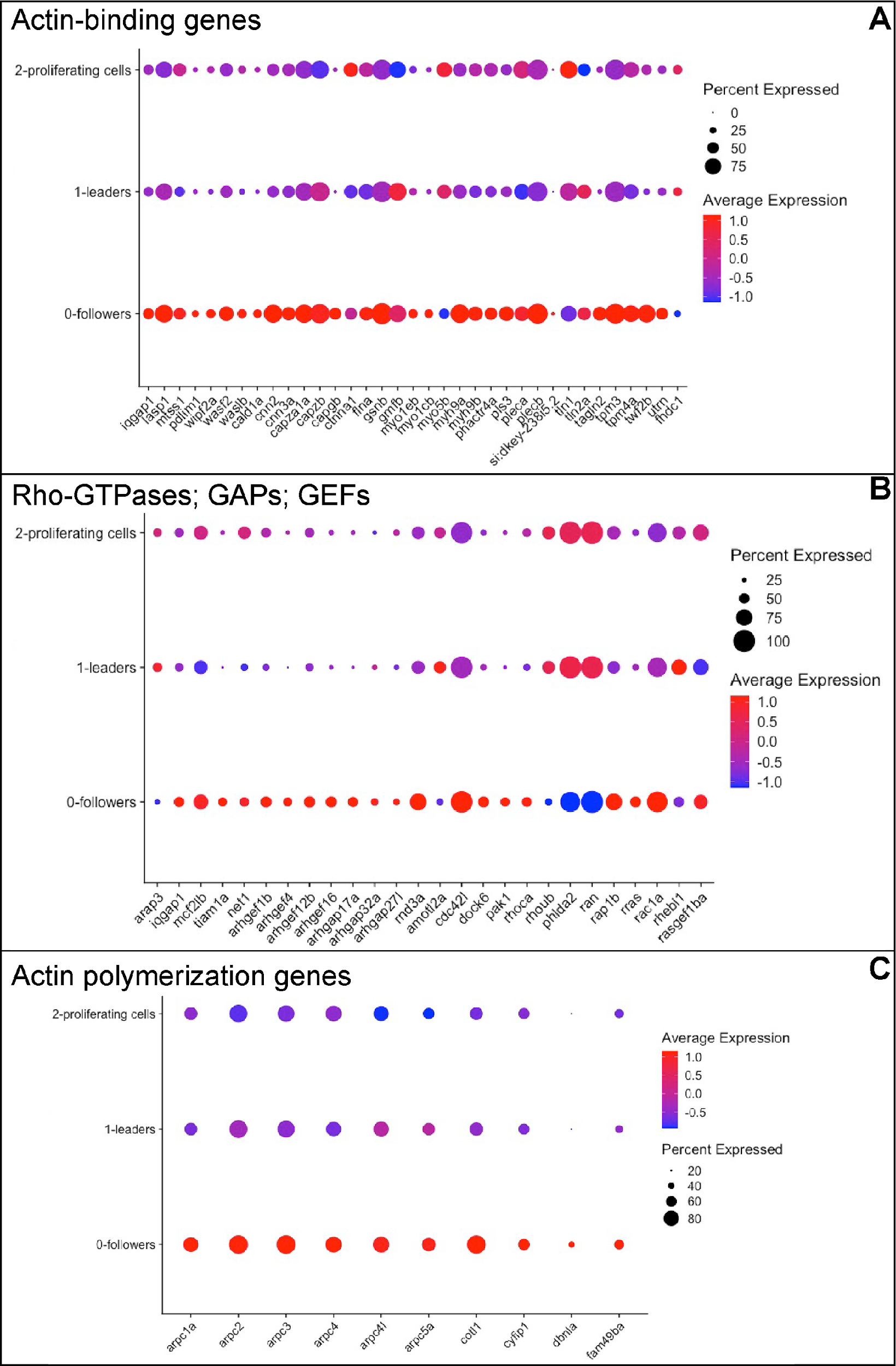

**Supplemental Figure 2: Expression of genes that regulate actin dynamics in pLLP clusters.**

(A-C) Using our scRNA-seq data set, we performed GO term enrichment analysis to reveal expression of actin-binding genes (A), Rho GTPases, GAPS, and GEFs (B), and actin polymerization genes (C) among the three pLLP specific clusters. Note enhancement of expression of actin-binding genes and actin polymerization genes in the follower population compared to the leaders and proliferating cells.

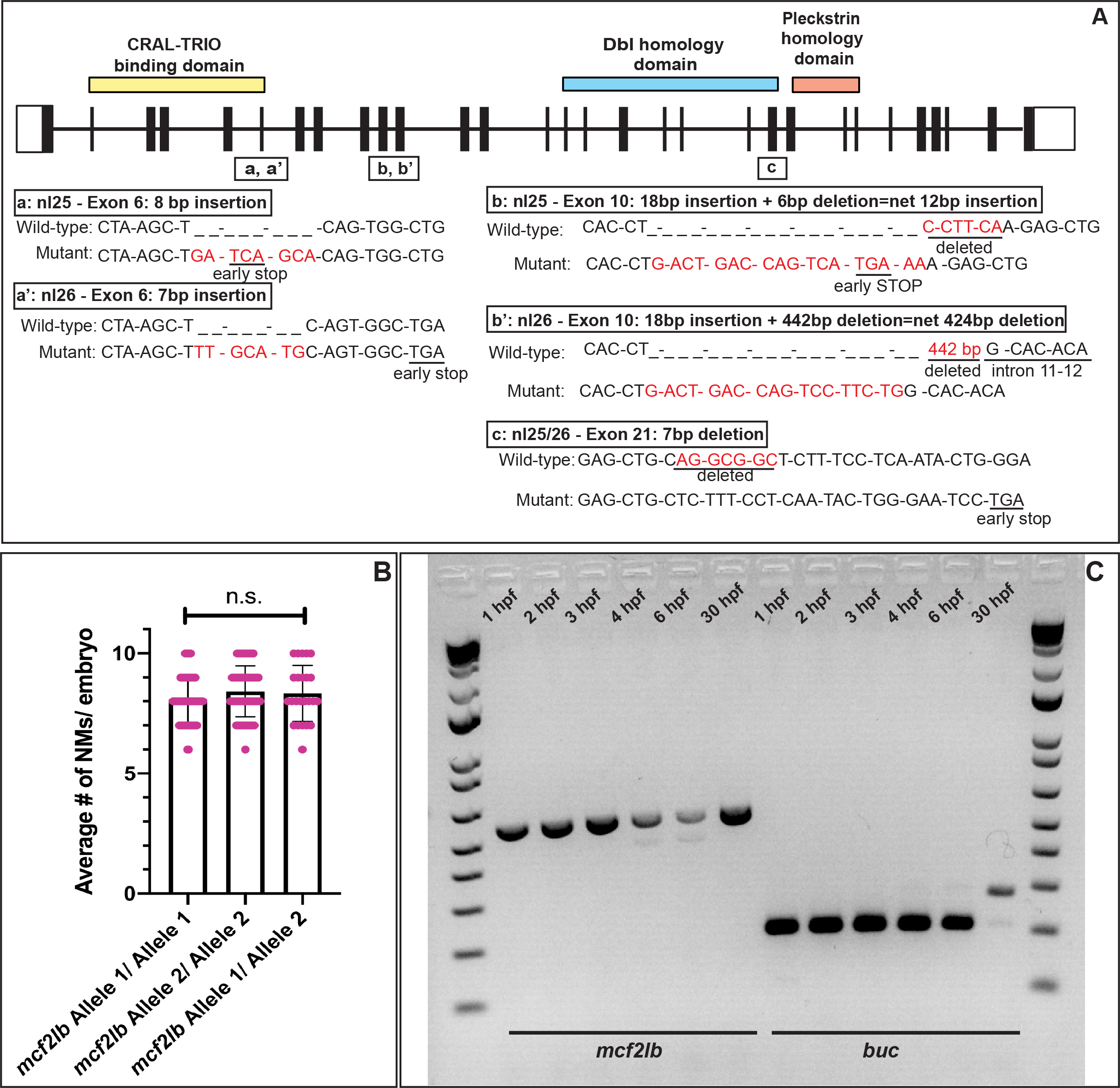

**Supplemental Figure 3: *mcf2lb* mutant validation.**

(A) Schematic and sequence of the four different CRISPR/Cas9-induced mutations. There are 2 mutations in exon 6, 2 mutation in exon 10, and 1 mutation in exon 21. This resulted in two alleles further designated as nl25 (Allele 1), nl26 (Allele 2). All mutations lead to early STOPs as indicated. (B) Phenotypic comparison between *mcf2lb* Allele 1/ Allele 1 (n = 56 embryos), *mcf2lb* Allele 2/ Allele 2 (n = 60 embryos), *mcf2lb* Allele 1/ Allele 2 (n = 24 embryos) . Note there is no significant differences in phenotype between the two alleles (one-way ANOVA). (C) Expression of *mcf2lb* and *buc* during early development 1 – 6 hpf and at 30 hpf

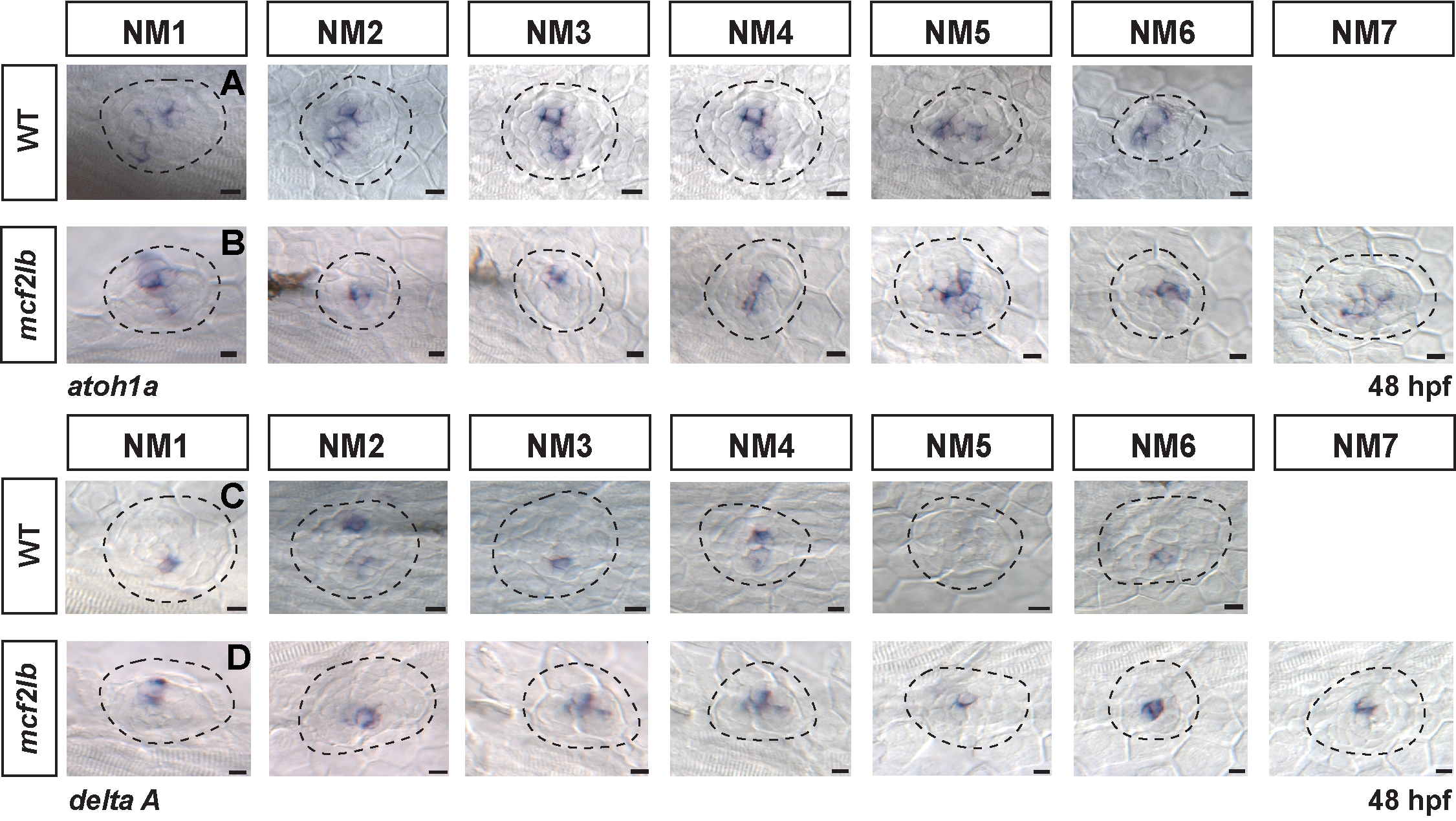

**Supplemental Figure 4: Hair cells are properly specified in the pLL of *mcf2lb* mutant embryos.**

(A – D) In situ hybridization of the hair cell markers *atoh1a* (A, B) and *deltaA* (C, D) in WT and *mcf2lb* mutant pLLs at 48 hpf. Dotted lines indicate NM. Scale bars = 5 μm.

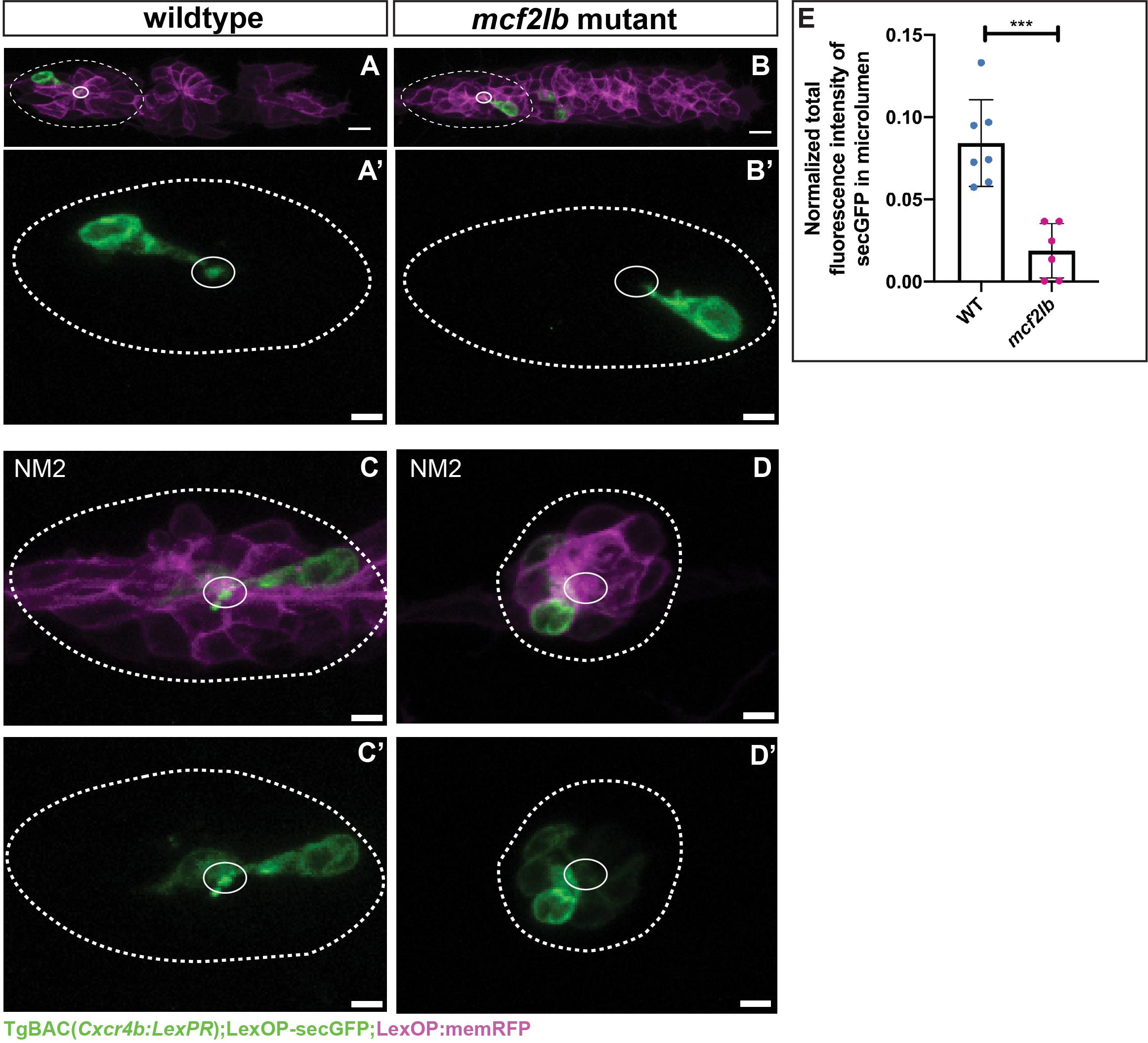

**Supplemental Figure 5: Microlumen intergrity is compromised in the *mcf2lb* mutant pLLP and differentiating NMs**.

LexOP-secGFP plasmid was injected into TgBAC(*Cxcr4b:LePR; LexOP:memRFP*) transgenic fish. LexPR was induced for 6 hours (29-35 hpf) to induce expression of secGFP (green) and memRFP (magenta). (A, B) secGFP expression in the trailing protoNM of WT and *mcf2lb* mutant pLLPs. Solid ovals denote presumptive microlumen and dotted line marks the trailing protoNM. (A’, B’) Higher magnidication of the protoNMs outlined in panels A and B. (C, D) secGFP expression in deposited NM2 of WT and *mcf2lb* mutant embryos (presumptive microlumen is outlined). (E) Quantification of fluorescence intensity of secGFP in the microlumen normalized to the fluorescence intesity of cells expressing secGFP that contribute to the rosette centers of trailing protoNMs and deposited NM2. *** p<0.001 (unpaired t-test). Scale bars in panels A, B= 10 μm and in panels A’, B’, C, C’, D, and D’ = 5 μm.

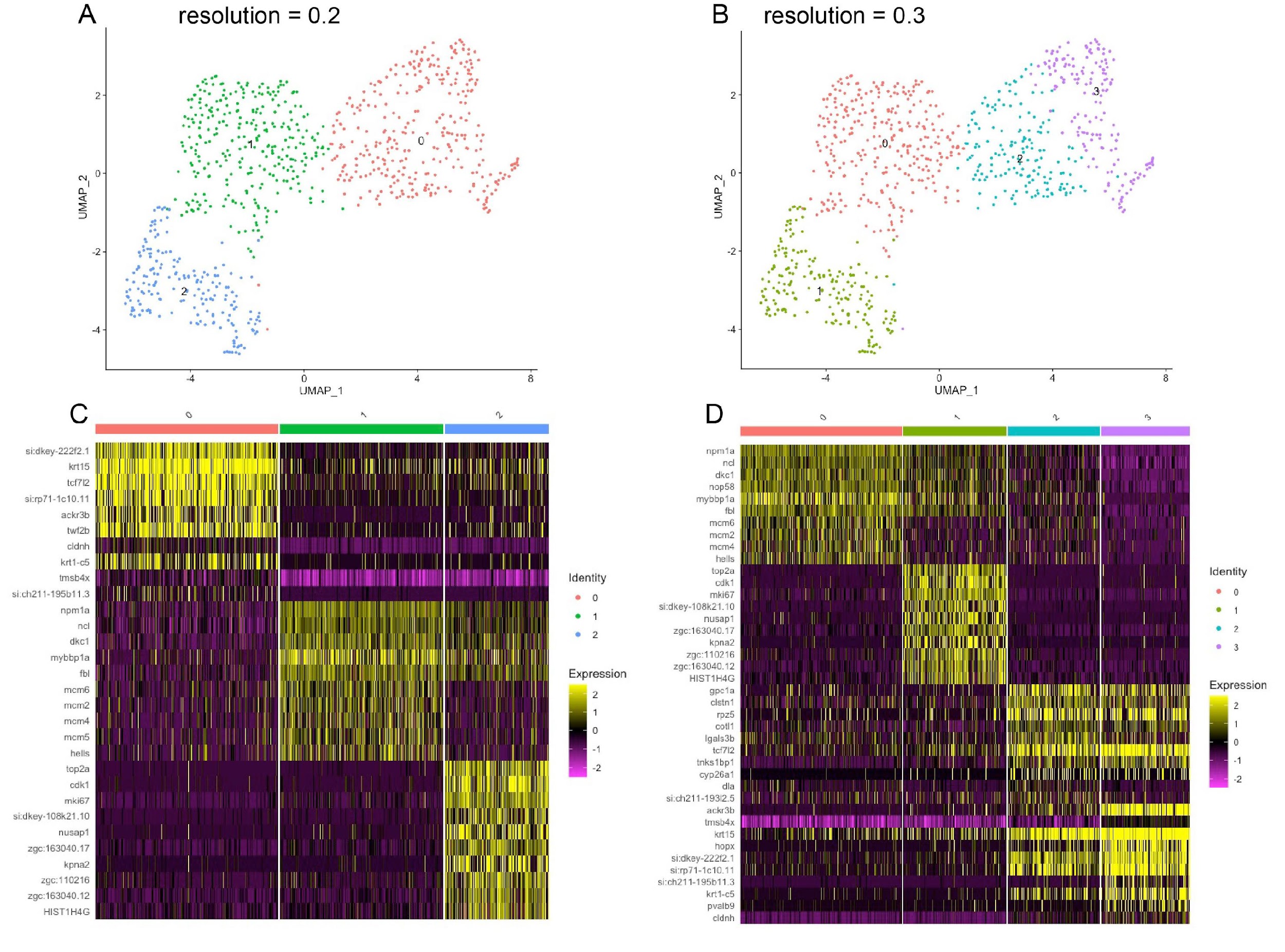

**Supplemental Figure 6. Subclustering of pLLP cells at different resolutions**.

A resolution sweep from 0.1 to 1 at 0.1 increments identified resolution 0.3 as optimally maximizing silhouette width. However, we noted that at this resolution, follower cells (cluster 0 in A) separated into two clusters that have extremely similar gene signatures (cluster 2 and 3 in B): compare gene signature for clusters 2 and 3 (D). This gene signature also contained 9 out of 10 genes from cluster 0 (C).

**MOVIE LEGENDS:**

**Movie 1:** WT pLLP migration and NM deposition. Time lapse confocal projections of Tg(*-8.0claudinB:lynGFP*)*^zf106^* expressing cells in a WT embryo. Embryo was imaged continuously starting at 30 hpf for 15 hours. Scale bar = 20 μm.

**Movie 2:** *mcf2lb* mutants show abnormal NM deposition behavior. Time lapse confocal projections of Tg(*-8.0claudinB:lynGFP*)*^zf106^* expressing cells in a *mcf2lb* mutant embryo. Embryo was imaged continuously starting at 30 hpf for 14 hours. Scale bar = 20 μm.

**Movie 3:** WT pLLP organization during migration. Time lapse confocal projections of Tg(*-8.0claudinB:lynGFP*)*^zf106^* expressing cells in a WT embryo at high magnification. Embryo was imaged continuously starting at 30 hpf for 1.5 hours. Scale bar = 10 μm.

**Movie 4:** *mcf2lb* mutants show abnormal NM deposition behavior. Time lapse confocal projections of Tg(*-8.0claudinB:lynGFP*)*^zf106^* expressing cells in a *mcf2lb* mutant embryo at high magnification. Embryo was imaged continuously starting at 30 hpf for 1.5 hours. Scale bar = 10 μm.

**Movie 5:** WT cells maintain contact with rosette center during pLLP migration. Time lapse confocal projections of a mosaically labeled WT embryo with WT Tg(*prim:lyn2-mCherry*) positive cells. Embryo was imaged continuously starting at 30 hpf for 1-1.5 hours. Scale bar = 10 μm.

**Movie 6:** *mcf2lb* mutant cells fail to maintain contact with rosette center during pLLP migration. Time lapse confocal projections of a mosaically labeled *mcf2lb* embryo with the *mcf2lb* Tg(*prim:lyn2-mCherry*) positive cells. Embryo was imaged continuously starting at 30 hpf for 1-1.5 hours. Scale bar = 10 μm.
